# Intracellular pH modulates vimentin remodeling in response to oxidants

**DOI:** 10.1101/2023.12.21.572888

**Authors:** Alma E. Martínez, Paula Martínez-Cenalmor, Patricia González-Jiménez, Cristina Vidal-Verdú, María A. Pajares, Dolores Pérez-Sala

## Abstract

Vimentin plays key roles in cell mechanosensing, cytoskeletal crosstalk and stress responses, and is finely tuned by posttranslational modifications. The vimentin single cysteine residue, C328, is a hotspot for modification and essential for filament remodeling by oxidants and electrophiles. With a pKa near physiological pH, C328 reactivity could be sensitive to cellular pH fluctuations. Here, we show that C328 modifications and vimentin reorganization by various reactive agents are modulated in response to pH variations. Lowering intracellular pH prevents, whereas intracellular alkalinization potentiates vimentin network disruption by oxidative and electrophilic species, including diamide, hydrogen peroxide and hydroxynonenal. The protective effect associated with low pH is selective for vimentin since it does not preclude oxidant-elicited disruption of actin or tubulin structures. Vimentin C328A and C328H mutants are resistant to disruption under all pH conditions, which highlights the importance of the thiol group at this position for the sensitizing effect of alkaline pH. Chemogenetic and optogenetic modulation of cellular pH allow spatiotemporal tuning of vimentin susceptibility to oxidants, suggesting the potential role of pH in the regulation of vimentin organization at precise locations. Alkalinization and generation of reactive oxygen species cooperate at the cell front during migration. We show that vimentin disassembly at cell edges of migrating fibroblasts, and lamellipodia formation, are affected by pH changes and the presence of C328. We propose that vimentin C328 could behave as a coincidental pH and redox responsive element, contributing to the precise regulation of vimentin assembly by the concert of these factors, illustrating the pH dependence of cysteine-mediated redox signaling.

## Introduction

The intermediate filament protein vimentin plays key roles in cell architecture and mechanotransduction [1, 2], and is also critical for organelle homeostasis, integration of signaling pathways and cellular responses to stress [3–5]. Indeed, although vimentin may seem dispensable for some processes under basal conditions, its broad implications are revealed under stress. Thus, vimentin-deficient mice show compromised wound healing, smaller size, vascular dysfunction and altered responses to infection, among other phenotypes [5].

Vimentin assembly into filaments is a complex process not completely elucidated. Most models support the formation of parallel dimers, which then form staggered antiparallel tetramers; these subsequently associate laterally to form “unit length filaments” (ULFs), which connect end to end to form filaments [6–9]. Vimentin filaments are highly dynamic structures that can elongate by addition of subunits at either end, exchange subunits at any point along their length, severe into fragments and fuse again in cells [10, 11]. This dynamic behavior is tightly modulated by posttranslational modifications (PTMs), and is required for efficient reorganization during essential cellular processes, such as mitosis and migration, as well as in response to stresses, including oxidative stress [12].

Indeed, oxidants and electrophiles elicit a profound, structure-dependent, remodeling of the vimentin network [12, 13]. Importantly, the single cysteine residue of vimentin, C328, is a key target for numerous PTMs, and is necessary for vimentin reorganization under oxidative or electrophilic stress [3, 14]. Structurally different C328 PTMs display non-overlapping functional consequences [15–17] (reviewed in [18, 19]). Modifications by certain small moieties, such as nitrosylation, have mild consequences on filament assembly, whereas bulkier moieties can even elicit filament severing [17]. Importantly, introduction of different side chains at the 328 position through mutagenesis elicits distinct morphological network rearrangements, which reinforces the structure dependence and functional significance of local perturbations at this site [13]. Interestingly, certain C328 mutants, including C328A and C328H form extended filament networks in cells, similar to wt, although they do not possess the thiol chemistry and therefore they are markedly resistant to modification or reorganization by thiol reactive agents [13].

The modifications of cysteine residues are central to redox signaling and oxidative stress responses [20–22]. Reversible cysteine modifications, such as sulfenylation, nitrosylation or glutathionylation, provide fast and flexible regulation of signaling pathways, and can constitute protective mechanisms avoiding excessive oxidative damage. Conversely, restoration of protein function after more stable modifications, such as sulfonylation or certain kinds of lipoxidation, may require new protein synthesis. Cysteine residues with catalytical or redox sensing functions often display a low pKa, which contributes to the presence of the thiol residue in the more reactive thiolate form [23]. Nevertheless, although pH-dependent modification of various protein cysteine residues has been reported in vitro [24, 25], demonstration of the pH-dependence of cysteine modifications in intact cells remains elusive. We have recently determined that C328 in purified human vimentin displays two ionization curves yielding pKas of approximately 4.5 and 7.4, suggesting the existence of two cysteine populations [26]. This implies that near physiological pH (7.2-7.4) a significant proportion of C328 residues (>50%) would be in the thiolate form [27]. Therefore, if a similar ionization behavior takes place in cells, changes in intracellular pH (pHi) in the physiological range could induce significant differences in C328 reactivity, and therefore in its extent of modification and subsequent impact on vimentin reorganization.

pHi is subjected to broad variations with important functional consequences, both in physiological and pathophysiological conditions, as well as in response to drugs. A pHi of approximately 7.2 is considered normal for cells in culture, whereas tumor cells display a more alkaline pHi (between 7.3 and 7.6) [28], which affords a proliferative and invasive advantage [29, 30]. Indeed, mitogens increase pHi by 0.1 to 0.3 pH units [31], and malignant transformation by certain oncogenes is accompanied by a rise in pHi [32]. In contrast, apoptosis by growth factor deprivation may associate with a marked intracellular acidification (pHi<6.8, [33]). Similarly, intense acidifications occur in muscle cells during high intensity exercise (pHi <6.8 [34]), as well as in certain models of hypoxia or ischemia [35, 36]. Lastly, temporal pHi variations occur within single cells depending on the cell cycle [37, 38], whereas spatial pHi variations include those between distinct organelles, from acidic lysosomes to the basic mitochondrial matrix [39, 40], as well as the intracellular gradients generated during cell migration [39, 41].

Here, we have monitored the effect of pH variations on vimentin C328 modifications and network remodeling by oxidants and electrophiles. Our results support that vimentin responsiveness to oxidative and electrophilic stress is strongly dependent on environmental pH, through mechanisms requiring the C328 thiol group. These observations unveil a previously underestimated factor contributing to the regulation of this intermediate filament protein and provide a relevant example of regulation of protein function through pH modulation of cysteine reactivity.

## Materials and Methods

### Reagents

HEPES, Pipes, urea, 2-mercaptoethanol, 5-(N-ethyl-N-isopropyl)amiloride (EIPA), syrosingopine and diamide were from Merck; Tris was obtained from VWR; DTT, pHrodo™ Red, and EZ-Link™ Maleimide-PEG2-Biotin (Mal-B) were from Thermo Fisher Scientific; Bis-carboxyethyl-carboxyfluorescein, acetoxymethyl ester (BCECF AM) was from Molecular Probes. H_2_O_2_ was from ChemCruz; 4-hydroxy Nonenal (HNE) was from Cayman Chemical. Precision Plus Protein Dual Color Standards and Sypro Ruby were from BioRad. PureCube Maleimide-Activated Agarose beads were from Cube Biotech. Anti-vimentin V9 (sc-6260) in its unconjugated, Alexa Fluor 488 and agarose conjugated versions was from Santa Cruz Biotechnology. Anti-vimentin D21H3 was from Cell Signaling Technology. Rabbit anti-α-tubulin antibody (ab52866) was from Abcam. Rabbit anti-GFAP was from DAKO. Alexa Fluor 647 anti-rabbit (A-21244) was from Thermo Fisher Scientific. Mouse TrueBlot ULTRA secondary antibody was from Rockland Immunochemicals. Immobilon-P membranes were from Millipore.

### Protein preparation

Recombinant human vimentin wild type (wt), and C328S, purified essentially as described [42] were obtained from Biomedal (Spain) or from Abvance (Spain). The protein in 8 M urea, 5 mM Tris-HCl pH 7.6, 1 mM EDTA, 10 mM 2– mercaptoethanol, 0.4 mM PMSF and approximately 0.2 M KCl was ultrafiltrated using Millipore Amicon Ultra filter units (10 K pore size) and dialyzed step-wise in 5 mM Pipes-Na pH 7.0, 1 mM DTT containing decreasing urea concentrations (8, 6, and 2 M) at room temperature. Final dialysis was performed for 16 h at 16 °C in 5 mM Pipes-Na pH 7.0 and 0.25 mM DTT [43]. The ultrafiltration step was necessary for efficient removal of EDTA [43]. Protein concentration was estimated from its A_280_, using an extinction coefficient of 22450M^−1^ cm^−1^, and buffer from the last dialysis step as blank. Refolded protein was subsequently subjected to DTT removal or stored directly in aliquots at –80 °C until use. For DTT elimination, vimentin underwent three cycles of 1:20 (v/v) dilution with 5 mM Pipes-Na pH 7.0 and ultrafiltration on Amicon Ultra filter units in order to reach a theoretical DTT concentration of 30 nM.

### Vimentin modification by reactive agents at several pH

The in vitro pH-dependent modification of purified and refolded vimentin was assessed by gel-based techniques, essentially as previously reported [12]. Briefly, vimentin at 4 µM final concentration, in 5 mM Hepes buffer at different pH (7.0, 7.5 and 8.0), was incubated at 24 °C in the presence of 1 mM H_2_O_2_ or 1 mM diamide for 1 h, or with 20 µM Mal-B for 15 min. The buffers at different pH used were near the pK_a_ values previously calculated by us [26]. Given the fact that DTT can react with electrophilic compounds and chelate metals [44], reactions of Mal-B with vimentin were carried out after DTT elimination. Additionally, Mal-B was added for a short incubation time in order to prevent its degradation, since the hydrolysis rate of this compound increases with pH and temperature [45]. In contrast, diamide shows a good oxidizing capacity over a broad range of pH [46]. For modifications with diamide and H_2_O_2_, the final DTT concentration was kept below 0.1 mM.

Reactions were stopped by boiling in Laemmli sample buffer in the absence (non-reducing conditions), or in the presence of 5% (v/v) 2-mercaptoethanol (reducing conditions), and proteins were analyzed by SDS-PAGE on 10% (w/v) polyacrylamide gels. Then, proteins incubated with Mal-B were electrotransferred to Immobilon-P membranes using a Trans-blot SD semi-dry transfer cell (Bio-Rad). Biotin incorporation was assessed by incubating blots with HRP-streptavidin (1:1000 v/v, GE Healthcare) and vimentin levels were estimated by western blot as described below. On the other hand, vimentin oligomerization after treatment with H_2_O_2_ or diamide was analyzed by SDS-PAGE followed by protein staining. Briefly, gels were fixed with 40% (v/v) methanol and 10% (v/v) acetic acid for at least 1 h, stained with Sypro Ruby from 90 min to 3 h, washed with 10% (v/v) methanol and 6% (v/v) acetic acid, and visualized under UV light on a Gel-Doc XR Imaging System (Bio-Rad). All the aforementioned steps were performed under gentle agitation and, importantly, the gel was protected from light as of the staining step. Signal intensities were evaluated by image scanning and analysis with ScionImage.

### Optical microscopy of purified vimentin structures

Vimentin at 4 µM was incubated in 10 mM HEPES pH 7.0 or 8.0, 0.1 mM DTT, in the absence or presence of 1 mM diamide for 1 h at room temperature. At the end of the treatment, vimentin polymerization was elicited by addition of 150 mM NaCl, final concentration, and the incubation was continued for 1 h at room temperature. Subsequently, anti-vimentin V9 antibody (1:200 v/v) was added for 30 min, followed by anti-mouse-Alexa Fluor 488 (1:200 v/v) for an additional 30 min. Aliquots from the incubation mixtures were deposited on coverslips, allowed to dry and mounted on glass slides with Fluorsave. Vimentin structures were visualized by fluorescence microscopy on a Leica SP5 microscope.

### Cell culture and treatments

SW13/cl.2 vimentin-deficient cells were the generous gift of Dr. A. Sarriá (University of Zaragoza, Spain) [47]. HeLa and U-251 MG cells were authenticated by microsatellite amplification (short tandem repeat (STR)-PCR profiling) at Secugen, S.L. (Madrid, Spain). Primary human dermal fibroblasts (ref. GM00409) were obtained from the NIGMS Human Genetic Cell Repository at the Coriell Institute for Medical Research (Camden, NJ). Mouse embryonic fibroblasts (MEF) from wild type (MEF wt) and vimentin knockout mice (MEF *Vim(-/-)*) were the generous gift of Prof. John Eriksson (Abo Academy University, Turku, Finland). They were cultured in DMEM with 10% (v/v) fetal bovine serum (FBS, from Sigma or Gibco) and antibiotics (100 U/ml penicillin and 100 μg/ml streptomycin). All cells were routinely tested for mycoplasma contamination. Treatments were generally performed in serum-free medium. Unless stated otherwise, cells were treated with 1 mM diamide for 15 min, either directly of after preincubation with 25 µM EIPA for 15 min, 10 µM monensin for 5 min, 10 µM syrosingopine for 1 h. When indicated, cells were treated with 10 µM HNE for 2 h or with 1 mM H_2_O_2_ for 25 min, after preincubation with EIPA or monensin as stated above. For treatment in acidic or in alkaline medium, 12 mM HCl or 6.25 mM NaOH, final concentrations, respectively, were added to DMEM containing 10 µM nigericin, and the cell culture medium was replaced with this mixture immediately before the addition of diamide. For some experiments, cells were seeded on 6-well plates containing one coverslip at the bottom of each well, as previously described [48]. After treatments, cells on coverslips were fixed and used for microscopy and cells attached to the wells were used for protein analysis, as described below.

### Plasmids and transfections

The RFP//vimentin wt, C328H and C328A bicistronic plasmids for expression of the red fluorescent protein DsRed-Express2 and vimentin as separate products, as well as the GFP-vimentin plasmid for expression of the fusion protein, have been previously described [3, 13]. mCherry-SEpHluorin was a gift from Prof. Sergio Grinstein (Addgene plasmid # 32001; http://n2t.net/addgene:32001; RRID:Addgene_32001) [49]. SW13/cl.2 cells were transfected with mCherry-SEpHluorin using Lipofectamine 2000 (Invitrogen), following the instructions of the manufacturer. The plasmid DsRed-stDCyD, bearing a nuclear export signal [50], was the generous gift of Prof. Emrah Eroglu and Dr. Ghaffari Zaki (Istanbul Medipol University). The plasmid YFP-NHE1 was obtained from Vectorbuilder. The plasmid pcDNA3.1-ArchT-BFP2-TSERex (ArchT-BFP), coding the light-gated outward proton pump ArchT fused to a blue fluorescent protein and an endoplasmic reticulum export sequence, was a gift from Yulong Li (Addgene plasmid # 123312; http://n2t.net/addgene:123312; RRID:Addgene_123312) [51]. SW13/cl.2 cells stably transfected with expression vectors for vimentin wt or C328A (RFP//vimentin wt or C328A), have been previously described [13].

### Cell lysis and western blot

Cells were washed with cold PBS and lysed in 50 mM Tris–HCl pH 7.5, 0.1 mM EDTA, 0.1 mM EGTA, 0.1 mM β-mercaptoethanol, containing 0.5% (w/v) SDS, 0.1 mM sodium orthovanadate and protease inhibitors (2 μg/ml each of leupeptin, aprotinin and trypsin inhibitor, and 1.3 mM Pefablock), by gentle scraping and forced passes through a 26 1/2G needle on ice. Protein concentration in lysates was determined by the bicinchoninic acid (BCA) method (ThermoFisher Scientific). Lysates were denatured at 95 °C for 5 min in Laemmli buffer and aliquots containing 30 μg of protein were separated on 10% (w/v) SDS-polyacrylamide gels. Proteins were transferred to Immobilon-P membranes (Millipore) using a semi-dry transfer system (Bio-Rad) and a three-buffer system following the instructions of the manufacturer. Blots were blocked with 2% (w/v) powdered skimmed milk in T-TBS (20 mM Tris-HCl, pH 7.5, 500 mM NaCl, 0.05% (v/v) Tween-20). Antibodies were diluted in 1% (w/v) BSA in T-TBS. Primary antibodies were routinely used at 1:1000 (v/v) dilution, followed by HRP-conjugated secondary antibodies (Dako) at 1:2000 (v/v) dilution. Bands of interest were visualized by enhanced chemiluminescence detection using the ECL system (Cytiva), exposing blots to X-Ray film (Agfa).

### Vimentin immunoprecipitation

SW13/cl.2 cells stably transfected with RFP//vimentin wt were incubated in control conditions or with diamide (1 mM final concentration for 15 min), and lysed as described above using a non-reducing lysis buffer (without β-mercaptoethanol) containing 1% (w/v) NP-40 instead of SDS, and 10 mM Mal-B for free thiol labelling. Lysates from control and diamide treated cells were then incubated for 30 min at room temperature. Aliquots of lysates containing 150 µg of protein were diluted with lysis buffer containing 1% (w/v) NP-40 to reach a final volume of 350 µl. Samples were then incubated overnight with anti-vimentin V9-agarose for immunoprecipitation at 4°C with gentle agitation. Beads were washed four times with cold lysis buffer containing 0.5% (w/v) SDS and 1% (w/v) NP-40, by centrifugation at 14,000 g for 1 min at 4°C, and subsequently boiled in Laemmli buffer during 5 min to elute the retained proteins. Eluted proteins were collected by centrifugation at 14,000 g for 2 min at room temperature. The immunoprecipitated products were separated by SDS-PAGE and transferred to Immobilon-P membranes. Mal-B incorporation was detected by incubation with HRP-streptavidin (1:6000 v/v), and the position of vimentin was confirmed by western blot using Mouse TrueBlot ULTRA as secondary antibody.

### Pull-down on maleimide-agarose

SW13/cl.2 cells stably transfected with RFP//vim wt were incubated with 1 mM diamide for 15 min, after preincubation with EIPA or monensin, as previously described. Cells were then washed twice with cold PBS and lysed in RIPA buffer containing protease inhibitors and, where indicated, 10 mM iodoacetamide. Lysates containing iodoacetamide were incubated for 30 min at room temperature to allow thiol alkylation. To minimize the presence of vimentin in oligomers that could lead to its indirect retention on the beads, 90 µg of total protein from cell lysates was diluted in urea (6.5 M final concentration). Samples were adjusted to the same final volume with coupling buffer I (100 mM NaH₂PO₄·2H₂O, 150 mM NaCl) and incubated overnight at 4°C with 80 µl of maleimide-activated agarose-conjugated beads. Agarose beads were pelleted by centrifugation at 500 g for 2 min at room temperature, and the supernatants (unbound proteins) were collected. Equal volumes of the unbound fractions from each condition were separated by SDS-PAGE and transferred to Immobilon-P membranes. Vimentin levels were detected by western blot using the anti-V9 antibody.

### Immunofluorescence and confocal microscopy

Cells on coverslips or on glass bottom dishes were washed twice with PBS and incubated with 4% (w/v) paraformaldehyde (PFA) in PBS for 25 min at room temperature. Fixed cells were permeabilized with 0.1% (v/v) Triton X-100 in PBS for 20 min at room temperature, and blocked with 1% (w/v) BSA in PBS for 1 h. This blocking solution was also used for antibody dilutions. For vimentin detection, cells were incubated with Alexa Fluor 488-conjugated anti-vimentin antibody (1:200 v/v) for 1 h, whereas for tubulin detection, anti-α-tubulin antibody (1:400 v/v) was used for 1 h followed by Alexa Fluor 647 conjugated secondary antibody (1:400 v/v) for an additional hour. F-actin was detected by incubation of fixed and permeabilized cells with 0.25 μg/ml phalloidin tetramethylrhodamine B isothiocyanate (phalloidin-TRITC, Sigma) in blocking solution. Where indicated, nuclei were counterstained with 2.5 µg/ml 4’,6-diamidino-2-phenylindol (DAPI) for 1 h. Images were acquired on SP5 or SP8 Leica confocal microscopes, using 63× magnification, routinely every 0.5 μm, and either single sections or overall projections are shown, as indicated. Unless stated otherwise, scale bars shown on images are 20 μm.

### Photoactivation of ArchT-BFP

The protocol was based on those of Wu et al., 2019, and Donahue et al., 2021 [51, 52]. HeLa cells cultured on glass bottom dishes were transfected with ArchT-BFP and mCherry-SEpHluorin or GFP-vimentin, as indicated, and imaged 48 h later in a thermostatized chamber. Fields with ArchT-BFP positive or negative cells were selected under the microscope. Initial images of single sections of ArchT-BFP, GFP-vimentin and bright field were acquired with a zoom of 2x and the following parameters: ArchT-BFP, λ_ex_ 405 and λ_em_ 420-465 nm; GFP-vimentin, λ_ex_ 400 and λ_em_ 505-550; bright field with illumination with the 561 nm laser. A region of interest (ROI) was selected in the bright field image to achieve a zoom of 14x, and a pulse of 4 s of the 561 nm laser at 75% of potency (equivalent to 2.25 mW) was applied. Zoom was then switched to 2x, and single images were acquired in the order, ArchT-BFP, GFP-vimentin, bright field. This sequence was repeated 4-5 times for a total of 7 min. ArchT-BFP negative cells stimulated with the 561 nm laser, as above, or cells photoactivated with the 488 nm laser at 100% potency (equivalent to 1.7 mW) for 5 s, were used as controls.

### Detection of reactive oxygen species

Cells were incubated with 10 µM 2′,7′-dichlorofluorescein diacetate (DCF-DA) for 30 min at 37°C in DMEM without phenol red, and immediately subjected to the photoactivation protocol in a thermostatized chamber. Images were acquired on a SP5 microscope, using λ_ex_ 488 nm and λ_em_ 505-555 nm.

### Estimation of intracellular pH

Variations in pHi under the assay conditions were confirmed by various methods. When indicated, pHi was estimated by monitoring the fluorescence of SW13/cl.2 cells transfected with mCherry-SEpHluorin, a ratiometric pHi indicator protein, in which mCherry fluorescence is stable at several pH, whereas SEpHluorin fluorescence is very acid-labile, resulting in a decrease in green/red fluorescence ratio at acidic pH [49]. For pHi calculations, red and green fluorescence was acquired using λ_ex_ 561 nm and λ_em_ 572-635 nm for red, and λ_ex_ 488 nm and λ_em_ 505-555 nm, for green fluorescence. To ensure the pH responsiveness of the green/red ratios measured under the various experimental conditions, standard measurements were performed by determining the SEpHluorin and mCherry fluorescence intensities of mCherry-SEpHluorin-transfected cells incubated in a K^+^-rich buffer (25 mM HEPES, 145 mM KCl, 0.8 mM MgCl_2_, 1.8 mM CaCl_2_, 5.5 mM glucose) containing nigericin 10 µM, and adjusted to three different pH values (6.9, 7. 5, and 8.5), thus causing pHi to match the extracellular pH. For estimation of the pH change elicited by photoactivation of ArchT-BFP, only the fluorescence of SEpHluorin was considered, since photoactivation with the 561 nm laser bleaches the fluorescence of mCherry. Measurements were carried out either on a Leica SP8 microscope equipped with a CO_2_ and thermostatized chamber or on a Widefield Multidimensional Microscopy System Leica AF6000 LX for live cell imaging. In addition, pHrodo Red (Invitrogen) was used as an indicator of pHi, following the instructions of the manufacturer. Briefly, cells were washed once with live cell imaging solution (LCIS: 20 mM HEPES, pH 7.4, 140 mM NaCl, 25 mM KCl, 1.8 mM CaCl_2_, 1 mM MgCl_2_) and incubated with the pHrodo Red staining solution (pHrodo Red, diluted first 1:20 (v/v) in concentrated Power Load solution, and then 1:100 (v/v) in LCIS, to yield a 1:2000 (v/v) final dilution, at 37°C for 30 min. Then, cells were washed with DMEM and were cultured under the indicated experimental conditions. For assessment of pHi, live cells were imaged on a Widefield Multidimensional Microscopy System Leica AF6000 LX using the Texas Red filter. Calibration was performed by incubation in LCIS medium at different pH in the presence of nigericin, as stated above. In assays employing the plasmid pH-Control, pHi was monitored with the fluorescent indicator BCECF AM, essentially as previously described [53]. Briefly, HeLa cells were incubated with 1 µM BCECF AM for 30 min at 37°C in DMEM without serum. Cells were washed in this medium and immediately imaged in a thermostatized chamber in the Leica SP5 microscope, following the instructions of the manufacturer. pHi responsiveness was confirmed by varying extracellular pH in the presence of nigericin, as detailed above. Examples of calibration curves are shown in Supplementary Figure 1.

### Image analysis

For image analysis, Leica LasX software or Image J (FIJI) were used. Figures show single sections of images or overlay projections, as indicated in the corresponding figure legends. Non-filamentous vimentin from in vitro assays was quantified as the mean fluorescence intensity of regions of interest defined in areas devoid of filamentous structures, which contained background signal and/or amorphous protein accumulations. For estimation of vimentin filament disruption in cells, the “Analyze Particles” command of Image J was used to monitor the number of particles in the experimental conditions. Cell circularity and distance were measured with Image J. The area of lamellipodia was measured with LasX, and fluorescence intensity profiles along the segments of interest were obtained with Image J.

### Statistical analysis

All experiments were performed at least three times. The number of determinations for each experimental condition and/or the sample size are given in the corresponding figure legends. GraphPad Prism 8.0 software was used to perform the statistical analysis. Normality of data sets was evaluated. The non-parametric Kruskal-Wallis test, followed by the post hoc Dunńs multiple comparisons test were used for data sets that did not fit a normal distribution or when the type of distribution could not be determined. These data are represented as box blots, where, unless specified otherwise, the central mark is the median, the edges of the box are the 25^th^ and 75^th^ percentiles, and the whiskers extend to the most extreme data points. Significance level is established at p<0.05 and is given on the corresponding graph and/or in the figure legend. For comparison of two data sets the Mann-Whitney test was used. For data sets with normal distribution, results are shown as average values ± standard error of mean (SEM). In this case, comparisons of multiple data sets were performed by the one-way analysis of variance (ANOVA) followed by Tukey’s multiple comparison test. Comparisons between two data sets was performed by the Student’s t-test for unpaired samples. Statistically significant differences are indicated on the graphs and/or in the figure legends, as follows: *p<0.05, **p<0.01, ***p<0.001, ****p<0.0001.

## Results

### Oxidation and alkylation of vimentin C328 in vitro depend on pH

The single vimentin cysteine, C328 can be modified in vitro by oxidants and electrophiles. C328 in purified vimentin displays a relatively low pKa [26], implying that its reactivity could be modulated by pH variations in the physiological range, as schematized in Fig. 1a. Here, we have monitored the effect of several reactive agents on vimentin in vitro under various pH conditions. Oxidants such as H_2_O_2_ or diamide promote the formation of disulfide-bonded vimentin oligomers detectable by SDS-PAGE under non reducing conditions (Fig. 1b and c) [26]. Rising the pH of the incubation mixture in the 7.0-8.0 range augments oxidized oligomer formation (Fig. 1b, c). Moreover, C328 alkylation by Mal-B, estimated by the intensity of the biotin signal incorporated into vimentin, also increases from pH 7.0 to 8.0 (Fig. 1d), whereas no incorporation occurs in the vimentin C328S mutant (Supplementary Figure 2). Therefore, C328 oxidation and alkylation increase with pH within the 7.0-8.0 pH range.

**Figure 1.**
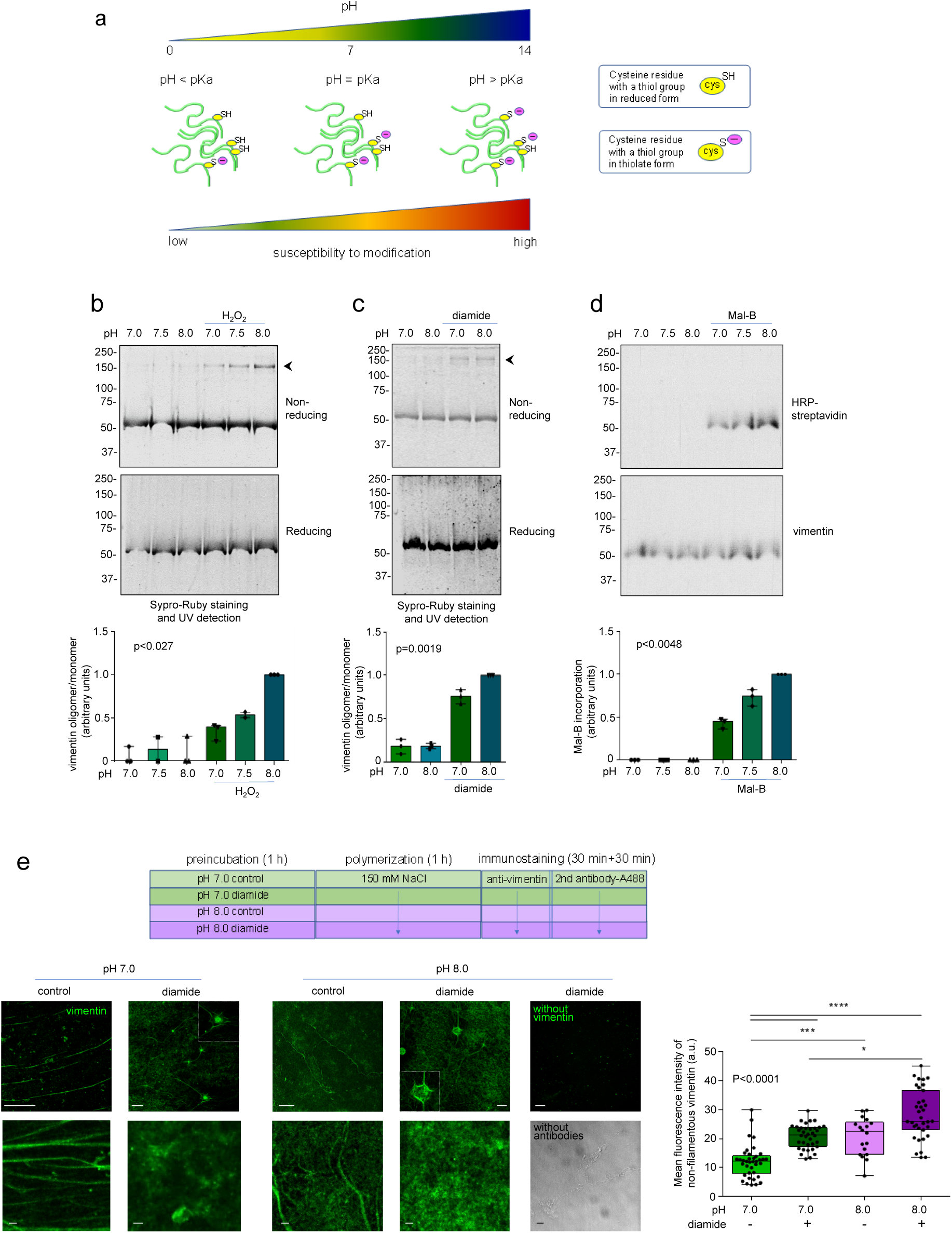
Modification of vimentin cysteine in vitro is sensitive to pH. (a) Scheme of the putative changes in cysteine ionization and modification depending on the pKa of the cysteine and the pH of the environment. (b-d) Modification of vimentin cysteine assessed by gel assays. Purified recombinant vimentin (4 µM) was incubated with various oxidants or cysteine reactive compounds, as follows: (b) 1 mM H_2_O_2_ for 1 h; (c) 1 mM diamide for 1 h or (d) 20 µM Mal-B for 15 min. (b, c) Aliquots from the incubation mixtures were analyzed by gel electrophoresis under non-reducing or reducing conditions, as indicated, and protein species were detected by total protein staining with Sypro Ruby. The position of the vimentin oligomers is indicated by an arrowhead (right), and that of molecular weight markers (kDa) is shown at the left. (d) Biotin incorporation was assessed by electrotransfer to Immobilon-P membranes and detection with HRP-streptavidin, whereas vimentin was detected with V9 anti-vimentin antibody. Graphs below the gels show box plots of the quantitation of vimentin oligomer vs monomer levels (b,c) or of biotinylated vimentin corrected by total vimentin (d). On each box, the top is the median, and the whiskers extend to the most extreme data points. Significance values obtained with the Kruskal-Wallis test are shown on graphs. (e) Purified vimentin was incubated in the absence or presence of diamide at different pH, as above, and subjected to polymerization with NaCl, as indicated in the assay outline (A488, Alexa Fluor 488). Subsequently, vimentin structures were visualized by immunodetection with the V9 antibody and fluorescence microscopy (scale bars represent 20 µm in upper images and 1 µm in lower images). Far right images show samples treated with diamide at pH 8.0, but without vimentin or without antibodies (bright field image), as indicated. At least three independent assays were performed and at least eight fields per assay were analyzed. The graph shows a box plot of the fluorescence intensity of non-filamentous vimentin, i.e. ROI devoid of filaments, displaying only background protein and aggregates, under the different experimental conditions, as detailed in Methods. On each box, the central mark is the median, the edges of the box are the 25^th^ and 75^th^ percentiles, and whiskers extend between the most extreme data points. Differences among conditions were found significant with the Kruskal-Wallis test (p < 0.0001), and between the indicated conditions by the post hoc Dunńs multiple comparisons test, *p<0.05, ***p<0.001, ****p<0.0001.

Vimentin polymerization in vitro yields filamentous structures detectable by fluorescence microscopy after immunolabeling [54]. Vimentin polymerized at pH 7.0 appears mainly as regular linear structures over a clean background with some small protein aggregates (Fig. 1e). Incubation with diamide before polymerization resulted in a higher background of disorganized protein, together with bright aggregates surrounded by some filamentous structures. Preincubation at pH 8.0 altered polymerization per se and potentiated disruption by diamide, further increasing background and eliciting large amorphous and condensed aggregates (Fig. 1e). Incubation mixtures treated in the same way, but excluding vimentin, only showed a dark background (Fig. 1e, upper right), whereas bright field observation of diamide-treated, non immunostained vimentin samples clearly showed the presence of aggregates (Fig. 1e, lower right). The fluorescence intensity of non-filamentous vimentin (regions devoid of filaments displaying background protein and aggregates) in the different experimental conditions is quantitated in Fig. 1e, graph. Together, these results show that changes in the pH of the assay mixtures modulate vimentin susceptibility to modification and disruption by oxidants in vitro.

### Remodeling of cellular vimentin by diamide depends on pHi

In cells, diamide induces an extensive remodeling of vimentin filaments into dots [3, 12, 13], which behave as biomolecular condensates [14]. Here, we have observed that diamide treatment progressively alters vimentin expressed in SW13/cl.2 cells, resulting in the appearance of bright dots after 15 min (Fig. 2a), which evolve towards a diffuse cytoplasmic background, evident after a 2 h treatment (Fig. 2a, and fluorescence intensity profile), indicative of filament disassembly and/or solubilization.

**Figure 2.**
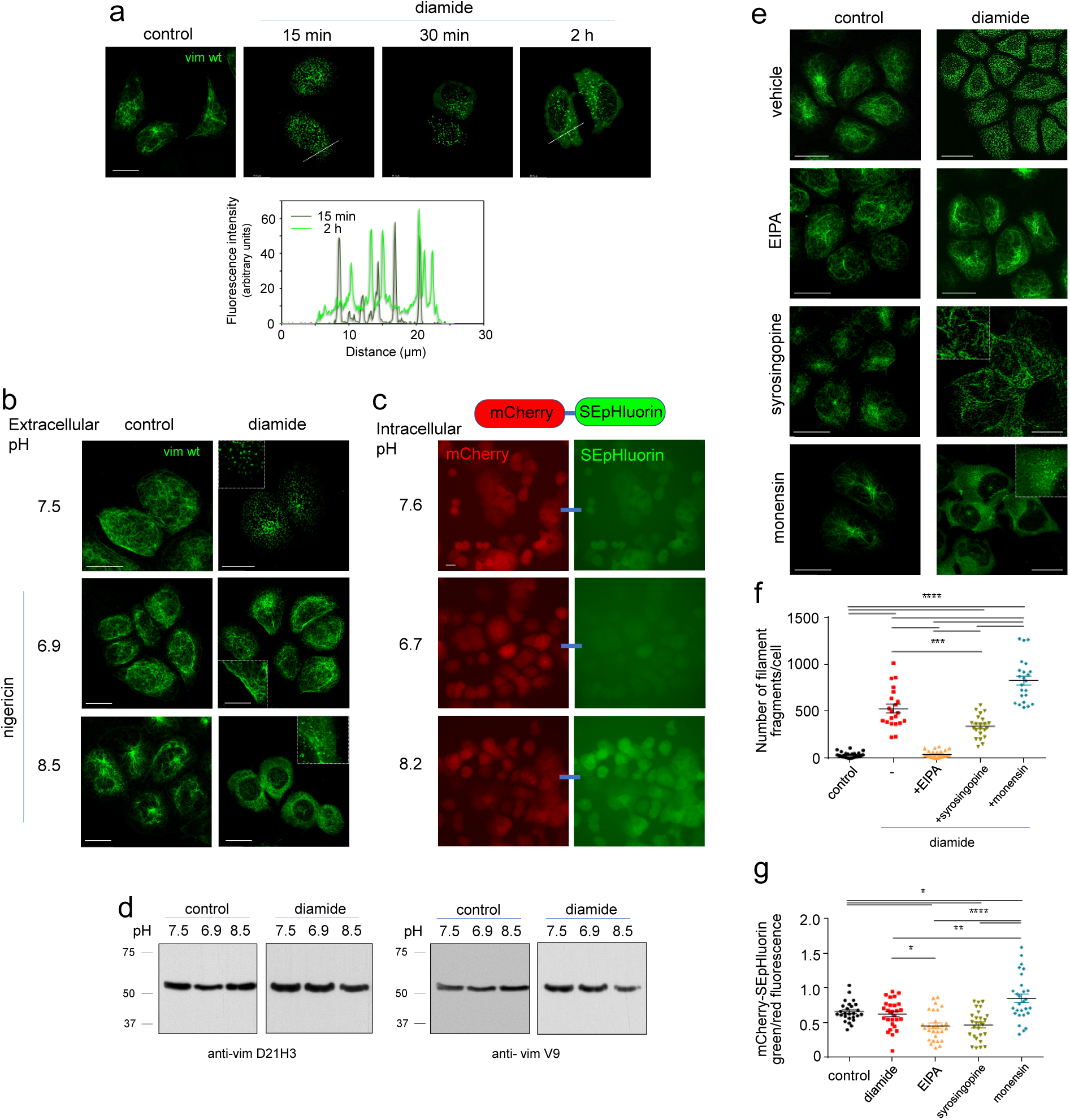
The disruption of the vimentin network by diamide is modulated by pH in cells. (a) SW13/cl.2 cells expressing vimentin wt were treated with 1 mM diamide for the indicated times and vimentin reorganization was assessed by immunofluorescence using V9 Alexa 488. Fluorescence intensity profiles along the dotted lines drawn on images from 15 min and 2 h treatments are depicted in the graph below illustrating the increase in cytoplasmic background related to vimentin disassembly. (b) Cells were incubated in culture media at different pH, supplemented with nigericin as indicated, and treated in the absence or presence of 1 mM diamide for 15 min. Vimentin was visualized by immunofluorescence as in (a). (c) SW13/cl.2 cells transfected with mCherry-SEpHluorin were incubated in media at different pH as in (b), and after 15 min, pHi was estimated from the ratio of the green/red fluorescence, as detailed in the experimental section (scale bars, 20 µm). (d) Lysates from SW13/cl.2 expressing vimentin wt treated as in (a) were analyzed by western blot with anti-vimentin antibodies directed to the N-terminus (D21H3, left images) or to the C-terminus (V9, right images). The position of molecular weight markers is shown at the left of blots. (e) SW13/cl.2 cells expressing vimentin wt were treated in the absence or presence of 1 mM diamide for 15 min after preincubation with 25 µM EIPA for 15 min, 10 µM syrosingopine for 1 h, or 10 µM monensin for 5 min and vimentin reorganization was assessed by immunofluorescence, as above. Insets show enlarged areas of interest. All images are overall projections and scale bars shown are 20 µm. (f) The extent of reorganization was evaluated by counting the number of filament fragments/cell. Three independent experiments were carried out and at least 21 cells were evaluated per experimental condition. (g) SW13/cl.2 cells expressing mCherry-SEpHluorin were treated as in (e), and the ratio of green/red fluorescence was measured as an index of pHi variations. Twenty seven cells were evaluated. Data in (f, g) are shown as average values ± SEM, and comparisons were made by ANOVA followed by Tukey’s multiple comparison test, *p<0.05, **p<0.01, ****p<0.0001.

Interestingly, the effect of diamide was clearly modulated depending on pHi changes. Thus, the remodeling of vimentin induced by a 15 min diamide treatment at neutral pH was suppressed when cells were incubated in acidic medium (pH 6.9) supplemented with the proton ionophore nigericin, as evidenced by the preservation of intact filaments (Fig. 2b). Conversely, the effect of diamide was potentiated under alkaline conditions (medium at pH 8.5 plus nigericin), as reflected by a disassembly of vimentin similar to that observed after a 2 h diamide treatment at neutral pH (Fig. 2a). Remarkably, variations of pH in the absence of diamide did not impair filament integrity to any appreciable extent (Fig. 2b, left images). The pHi attained under the assay conditions was monitored in cells transfected with the ratiometric pHi indicator construct mCherry-SEpHluorin [49] (Fig. 2c).

A potential effect of the treatments on the integrity of the protein was assessed by western blot analysis of cell lysates obtained under the same experimental conditions. Detection of vimentin with antibodies directed towards the N-or the C-terminal domains of the protein (clones D21H3 and V9, with epitopes located around residues R45 and N417, respectively) evidenced the preservation of the full-length protein and the absence of degradation products (Fig. 2d). Thus, the marked remodeling of the vimentin network elicited by diamide at neutral and alkaline pHi is not associated with protein degradation, and could be attributed to filament disassembly.

### Vimentin disruption by diamide is modulated by inhibitors of ion transporters affecting pHi

pHi is subjected to a tight control involving numerous transporters, which can be modulated pharmacologically [31, 39] to elicit pHi variations. We observed that preincubation of cells with EIPA, which inhibits the Na^+^/H^+^ antiporter provoking intracellular acidification [53, 55], clearly protected the vimentin network from diamide-elicited disruption (Fig. 2e). Also, pretreatment with syrosingopine, which lowers pHi by inhibiting the lactate transporter [56], afforded a partial protection of filament integrity, with persistence of numerous filament remnants (Fig. 2e, inset).

Conversely, preincubation with the H^+^ ionophore monensin, which causes intracellular alkalinization [57], markedly potentiated the effect of diamide, leading to vimentin fragments together with a diffuse cytoplasmic signal indicative of vimentin disassembly (Fig. 2e), analogous to that observed by treatment with diamide in alkaline pH (shown in Fig. 2b). The severity of vimentin disruption was estimated by quantitation of the number of vimentin filament fragments/particles per cell (Fig. 2f). Diamide alone significantly increased the number of fragments with respect to control. EIPA pretreatment virtually blunted and syrosingopine pretreatment attenuated the effect of diamide, whereas monensin amplified its impact, as reflected by 1.5-fold increase in the average number of vimentin particles with respect to diamide per se (Fig. 2f). Of note, none of the compounds employed disrupted vimentin filament integrity on their own, although monensin induced a partial accumulation of filaments near the center of the cell (Fig. 2e). Monitoring mCherry-SEpHluoring fluorescence showed a decrease in the green/red fluorescence ratio with EIPA and syrosingopine and an increase with monensin, confirming that the pharmacological strategies employed modulated pHi in the directions expected (Fig. 2g).

### Variations in pHi affect vimentin redox responsiveness in various cell types

Next, we confirmed the modulatory effect of pHi in various cell types (Supplementary Figure 3). In HeLa epithelial cancer cells, diamide elicited an extensive disassembly of vimentin filaments into dots (Supplementary Figure 3a). This effect was attenuated by intracellular acidification through incubation at low extracellular pH in the presence of nigericin or by pretreatment with EIPA or syrosingopine. In sharp contrast, diamide elicited vimentin disruption was potentiated by rising pHi through treatment in alkaline medium plus nigericin or pretreatment with monensin, which led to diffuse disassembled vimentin coexisting with dots and some filament remnants. pHi variations elicited by the various strategies were confirmed with pHrodo (Supplementary Figure 3b). A similar modulatory effect of pHi was observed in primary human fibroblasts (Supplementary Figure 3c), in which diamide-provoked filament fragmentation was prevented by EIPA or the combination of nigericin and acidic medium. Conversely, incubation with diamide at alkaline pH or in combination with monensin led to a diffuse vimentin background, indicative of disassembly, in extensive areas of the cell (Supplementary Figure 3c). Thus, a modulatory effect of pHi on diamide-elicited vimentin disruption was evidenced in several cell types, all of which showed a protective effect of pHi acidification and a deteriorating effect of pHi alkalinization strategies.

We have recently reported that H_2_O_2_ and the electrophilic aldehyde hydroxynonenal (HNE) disrupt vimentin organization in cells by mechanisms dependent, at least in part, on the presence of C328 [13]. Interestingly, vimentin network disruption by these agents was also sensitive to pHi changes (Supplementary Figure 4). Incubation of SW13/cl.2 cells expressing vimentin wt with H_2_O_2_ induced vimentin bundles and/or juxtanuclear accumulations (Supplementary Figure 4a). These alterations were attenuated by treatment at acidic pH, where cells showed a more homogeneous filament distribution, and clearly potentiated at alkaline pH, where cells with filament fragments, dots, and/or diffuse vimentin were also apparent (Supplementary Figure 4a). HNE treatment elicited filament condensation into curly bundles and/or retraction from the cell periphery in MEF wt (Supplementary Figure 4b). These effects of HNE were mitigated by preincubation with EIPA, which favored a more extended network, and worsened by monensin, which potentiated filament condensation leading to the formation of aggresome-like structures (Supplementary Figure 4b). These observations indicate that the susceptibility of the cellular vimentin network to disruption by various oxidants and electrophiles depends on pHi.

### Intracellular acidification selectively protects vimentin against diamide elicited disruption

We next explored whether variations in pHi affect the susceptibility of other cytoskeletal structures to disruption by oxidants. In control SW13/cl.2 cells expressing vimentin wt, f-actin was mainly distributed at the cell cortex, as well as in short fibers of irregular orientation throughout the cytoplasm (Fig. 3a). In overall projections, the intensity of f-actin staining was rather homogeneous across the cell (Fig. 3a). Treatment with diamide elicited a characteristic f-actin pattern, consisting of a thin actin cortex and a cytoplasmic mesh of diffuse structures with a “sponge-like” pattern (Fig. 3a). Moreover, the intensity of f-actin staining decreased in the central region of the cell, as illustrated in the fluorescence intensity profiles of overall projections (Fig. 3a, graph). Remarkably, acidic conditions, elicited by incubation in medium at pH 6.9 in the presence of nigericin, did not prevent actin disruption by diamide (Fig. 3b), which also occurred upon treatment at alkaline pH. Acidic or alkaline pH conditions per se did not elicit a significant perturbation of the distribution of actin into small fibrils (Fig. 3b).

**Figure 3.**
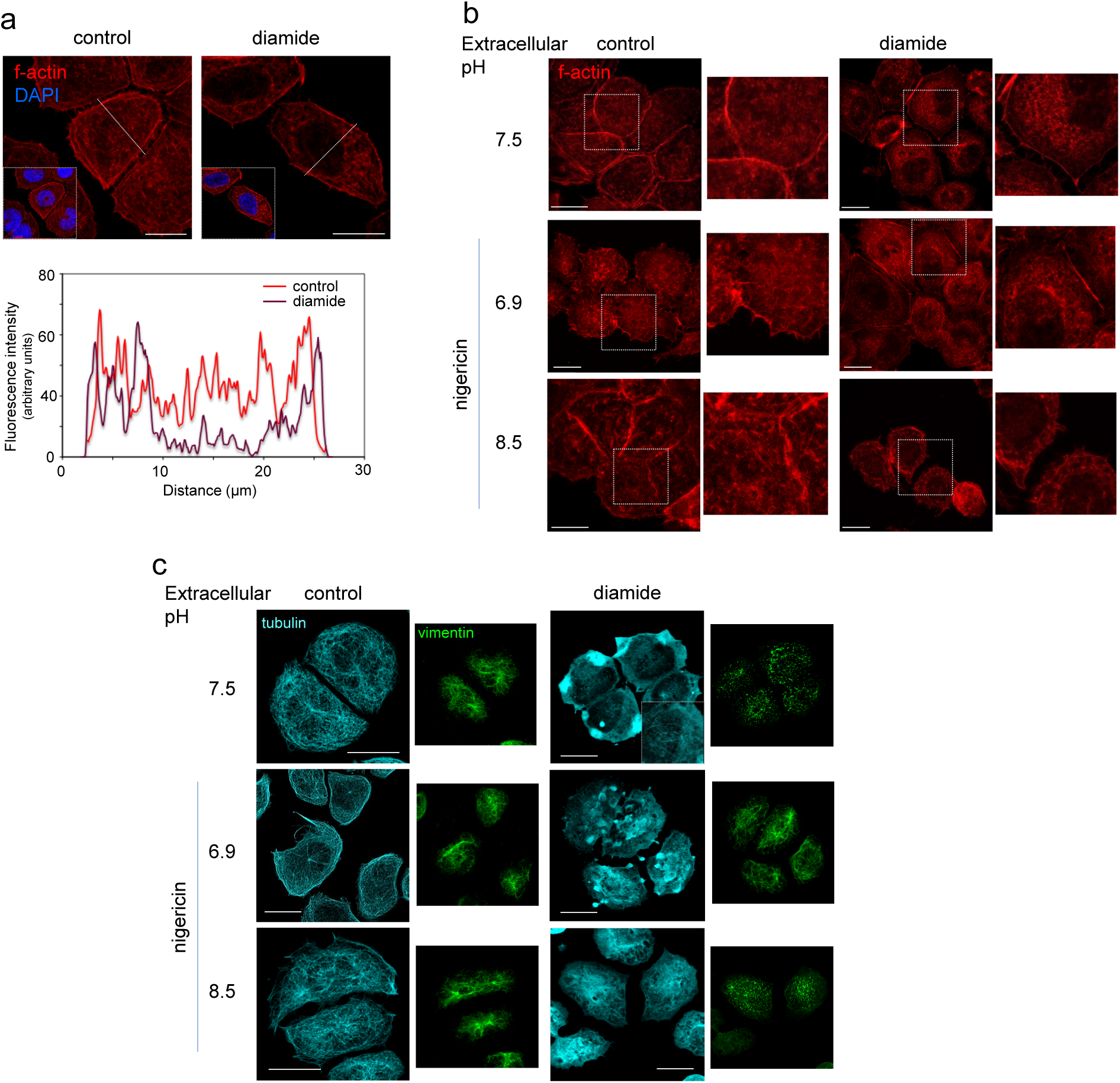
Effect of pHi variations on the disruption of other cytoskeletal structures by diamide. SW13/cl.2 cells expressing vimentin wt were incubated in culture media at various pH supplemented with nigericin, as indicated, and treated in the absence or presence of 1 mM diamide for 15 min. At the end of the treatment cells were fixed and stained with TRITC-phalloidin to visualize f-actin (a and b) or subjected to immunofluorescence with anti-tubulin and anti-vimentin antibodies (c). (a) The typical reorganization of f-actin elicited by diamide is illustrated. Insets show the position of nuclei stained with DAPI. Fluorescence intensity profiles along the dotted lines drawn on images are shown in the graph below to illustrate the decrease in f-actin staining coinciding with the nuclear region. (b) Representative images of actin alterations under the various experimental conditions are shown. Enlarged views of areas of interest delimited by dotted squares are depicted at the right of each image. (c) Representative images of tubulin alterations are shown (cyan) and the vimentin staining in the same cells is shown at the right of each image (green). All images are overall projections. Scale bars, 20 µm.

Diamide also profoundly altered microtubules, which lost their homogenous distribution and appeared as diffuse structures together with some condensations and scarce remaining misoriented microtubules (Fig. 3c, diamide, inset). pH variations did not elicit significant alterations of microtubules per se. Nevertheless, neither acidic nor alkaline pH conditions protected microtubules from diamide effects (Fig. 3c). Vimentin organization, shown here for comparison, illustrates that the preservation of vimentin filaments at acidic pH coexists with deeply perturbed microtubules. Thus, of the three main cytoskeletal networks, the modulatory effect of pHi variations on oxidant-elicited disruption is selective for vimentin.

Within intermediate filaments, proteins of the type III class, i.e., vimentin, glial fibrillary acidic protein (GFAP), desmin and peripherin, possess highly homologous sequences, and share a conserved cysteine residue, which, except in peripherin, is the only one in the monomer [19]. U-251 MG astrocytoma cells express GFAP, which is 58% identical to vimentin. Treatment of these cells with diamide induced the remodeling of GFAP filaments into fragments (Supplementary Figure 5). Notably, analogous to vimentin, this reorganization was also prevented by preincubation with EIPA, with conservation of filament continuity, and apparently worsened by monensin, thus showing that diamide-elicited remodeling of GFAP is also modulated by strategies altering pHi (Supplementary Figure 5).

### Variations in pHi influence vimentin C328 modification in cells

According to our hypothesis (Fig. 1a), the protective effect of acidic pHi on vimentin disruption by oxidants could be related to a decrease in the proportion of the thiol group of the single cysteine residue, C328, in its thiolate, reactive form, resulting in a lower susceptibility to chemical modification, as illustrated in vitro in Fig. 1b to d. Therefore, we assessed whether pHi variations influenced C328 modification elicited by diamide. For this, we have estimated the proportion of vimentin with a free thiol group as an inverse index of the extent of C328 modification, by using two complementary strategies (schematized in Fig. 4a). In the first approach (Fig. 4a, left branch), cells were treated in the absence or presence of diamide, and immediately collected and lysed in buffer containing Mal-B to label free thiol groups. Subsequent vimentin immunoprecipitation and biotin detection indicated that diamide treatment attenuated Mal-B incorporation into vimentin, suggesting that it diminishes free-thiol content (Fig. 4b). Additionally, strategies eliciting intracellular acidification or alkalinization by pretreatment with EIPA or monensin, respectively, tended to preserve (acidic pHi) or decrease (alkaline pHi) Mal-B incorporation with respect to the effect of diamide alone, suggesting that pHi variations influence vimentin thiol accessibility (Fig. 4c). In the second approach, urea was added to cell lysates at 6.5 M final concentration to dissociate vimentin oligomers. Then, proteins with free thiol groups were pulled down on maleimide conjugated beads (Fig. 4a, right branch). Vimentin from control samples was majoritarily retained on the beads, whereas vimentin present in lysates from cells treated with diamide or monensin plus diamide was clearly detected in the unbound fraction (Fig. 4d).

**Figure 4.**
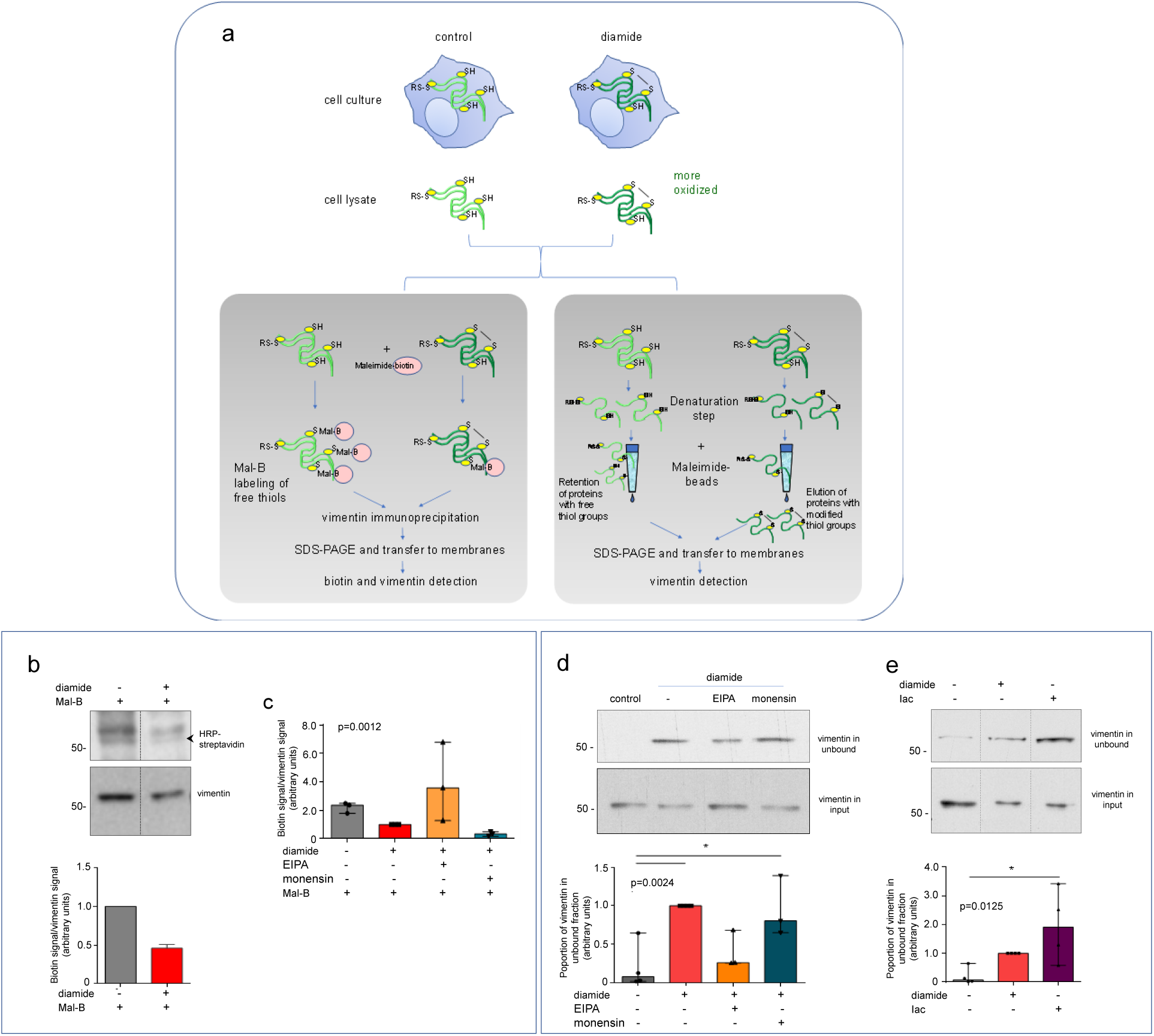
Effect of pHi modulation on the extent of modification of vimentin C328 in cells. (a) Scheme of the strategies employed to estimate the extent of modification of C328. (a, left branch) Cells treated with the compounds of interest, e.g., diamide, are lysed in the presence or absence of biotinylated maleimide (Mal-B) to label free thiols. Total lysates are subjected to vimentin immunoprecipitation, and immunoprecipitates analyzed for biotin and vimentin detection as detailed in Methods. High Mal-B incorporation is expected when cellular vimentin C328 is mainly in its free thiol form, whereas a low Mal-B signal indicates a higher degree of oxidation or modification under the treatment conditions. (a, right branch) Cell lysates are subjected to pull-down using maleimide conjugated agarose-conjugated. A higher vimentin abundance in the unbound fraction is expected when vimentin C328 is oxidized, whereas a lower signal indicates a higher proportion of C328 in free thiol form, and therefore greater retention on the maleimide conjugated beads. (b, c) SW13/cl.2 cells expressing vimentin wt were treated with 1 mM diamide for 15 min, or incubated with vehicle, 25 µM EIPA for 15 min or 10 µM monensin for 5 min before diamide treatment, as indicated, and processed according to the scheme in (a, left branch). The biotinylated polypeptide comigrating with vimentin is marked by an arrowhead. The dashed line indicates where lanes from the same gel have been cropped. Quantitation of biotin incorporation into the vimentin band, corrected by vimentin levels is shown in the graphs, expressed with respect to the control condition in (b) and with respect to diamide in (c). In (b), results from three independent assays are expressed as average values ± SEM. In (c), results from three assays are presented as box plot, where the top of each box in the median, and the whiskers extend from minimum to maximum values. Statistical significance was evaluated with the Kruskal-Wallis test (p value shown on the graph). (d, e) Lysates from SW13/cl.2 cells expressing vimentin and treated as above, were processed as schematized in (a, right branch). Vimentin was detected in the unbound (upper blot) and input (lower blot) fractions from pull-down on maleimide beads by western blot. (d) Proportion of vimentin in the unbound fraction from cells treated with diamide alone or after preincubation with EIPA or monensin, as indicated. (e) Comparison of the behavior of lysates from control and diamide treated cells, with that of a control lysate incubated with iodoacetamide (Iac) in vitro prior to pull-down. A minimum of three independent assays was performed for each approach. Results are shown as box plots, where the top of each box depicts the median value for every experimental condition, and the whiskers extend between the minimum and maximum values. Data sets were compared with the Kruskal-Wallis test (p values shown on the corresponding graphs), followed by the pos hoc Dunn’s test for multiple comparisons, *p<0.05.

In contrast, incubation of cells with EIPA before diamide treatment resulted in a proportion of vimentin in the unbound fraction closer to control lysates, suggesting a preservation of free thiol availability with respect to diamide alone (Fig. 4d). As a control for this assay, a cell lysate was treated with iodoacetamide prior to pull-down. This resulted in a greater recovery of vimentin in the unbound fraction, as expected due to C328 alkylation (Fig. 4e). Taken together, these results indicate modification of the C328 thiol upon treatment with diamide, which would be decreased by eliciting intracellular acidification with EIPA, thus suggesting that pHi impacts the oxidation of the vimentin thiol group.

### C328 is required for diamide elicited vimentin disruption at several pHi

We previously showed that C328 is required for the remodeling of the vimentin network by diamide [3, 12, 14]. To explore the importance of the C328 thiol group in the modulation of oxidative remodeling by pHi, we monitored the effect of diamide in MEF *Vim(-/-)* transfected with vimentin wt (Fig. 5a), or vimentin C328H or C328A mutants (Fig. 5b, c), in the absence or presence of inhibitors of ion transporters, namely, EIPA and monensin. These mutants were chosen on the basis of their ability to form regular vimentin networks in cells, and their resistance to thiol-modifying agents [3, 13, 14]. Whereas vimentin wt disruption by diamide was modulated by pharmacological alteration of pHi, vimentin C328H filaments were resistant to diamide-elicited disruption not only under control and acidic conditions, but also after pretreatment with monensin (Fig. 5b), a condition in which vimentin wt filaments are extensively disassembled (Fig. 5a). A virtually total protection was also afforded by the C328A mutation (Fig. 5c). The importance of C328 was confirmed in SW13/cl.2 cells transfected with vimentin wt or C328A (Supplementary Figure 6a). In this cell type, only vimentin wt was susceptible to filament remodeling by diamide, which was attenuated by EIPA and potentiated by monensin, whereas vimentin C328A formed filaments under all experimental conditions (Supplementary Figure 6a). Therefore, these observations suggest that C328, with its reactive thiol group, plays a key role not only in the remodeling of vimentin by oxidants, but also in the pHi-dependent modulation of this process.

**Figure 5.**
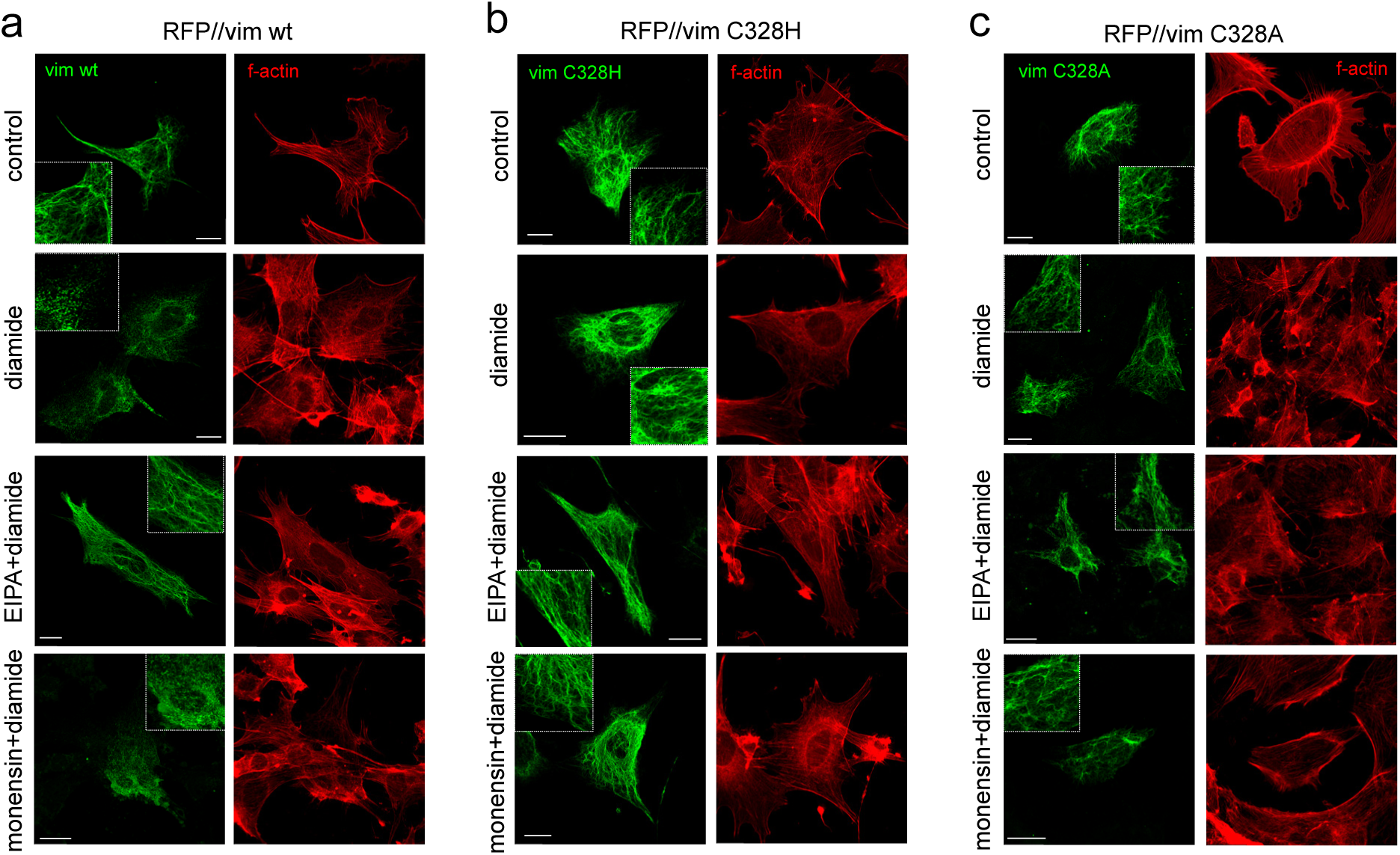
Importance of C328 in vimentin remodeling by diamide in the presence of pharmacological modulators of pHi. MEF *Vim(-/-)* were transfected with plasmids coding for vimentin wt (a) or the C328H (b) or C328A mutants (c). As indicated, cells were pretreated with vehicle, 25 µM EIPA for 15 min or with 10 µM monensin for 5 min, before treatment with 1 mM diamide for 15 min. The distribution of vimentin filaments was assessed by immunofluorescence (left images of each panel) and that of f-actin by staining with TRITC-phalloidin (right images of each panel). All images are overall projections; scale bars, 20 µm). Insets display enlarged areas of interest.

Interestingly, although vimentin C328H and C328A were protected from the effect of diamide, f-actin still displayed the typical decrease in f-actin staining at the center of the cell (Fig. 5a to c, right images, and Supplementary Figure 6b). These observations corroborate that substituting vimentin C328 by less or non-nucleophilic amino acids, selectively protects vimentin, but not other cytoskeletal structures, against the effect of diamide, and its modulation by pHi changes.

### Genetic pHi modulation allows spatiotemporal tuning of vimentin remodeling by oxidative stress

A chemogenetic approach (Fig. 6a), was used to elicit intracellular acidification in individual cells through transfection of the DsRed-stDCyD expression plasmid (pH-Control), which codes for the *Salmonella typhimurium* enzyme D-cysteine desulfhydrase (stDCyD) [50]. This enzyme converts the unnatural amino acid β-chloro-D-alanine (β-Cl-D-Ala) into its corresponding α-ketoacid, generating intracellular HCl as the main byproduct [50]. Here, we have confirmed that supplementing the cell culture medium with β-Cl-D-Ala selectively lowers pHi in HeLa cells transfected with a cytoplasm targeted pH-Control vector (Fig. 6b). Consistently, β-Cl-D-Ala protects the vimentin network from oxidative stress induced by diamide selectively in cells expressing pH-Control (Fig. 6c). In the absence of β-Cl-D-Ala, diamide elicits extensive conversion of filaments into dots in both pH-Control positive and negative cells (Fig. 6c), whereas in the presence of the β-Cl-D-Ala substrate, filament integrity is preserved particularly in pH-Control transfected cells. These effects are reflected in the proportion of cells displaying vimentin dots under the different experimental conditions (Fig. 6c, graph). This strategy also allows temporal regulation of pHi [50]. Therefore, we next tested whether removing β-Cl-D-Ala from the incubation medium abolished the protective effect. Indeed, whereas incubation of pH-Control positive cells with diamide plus β-Cl-D-Ala for 30 min afforded filament protection, removal of β-Cl-D-Ala after the first 15 min of treatment, unleashed filament disruption (Fig. 6d), thus indicating that vimentin remodeling by diamide can be reversibly modulated by pHi at the single cell level.

**Figure 6.**
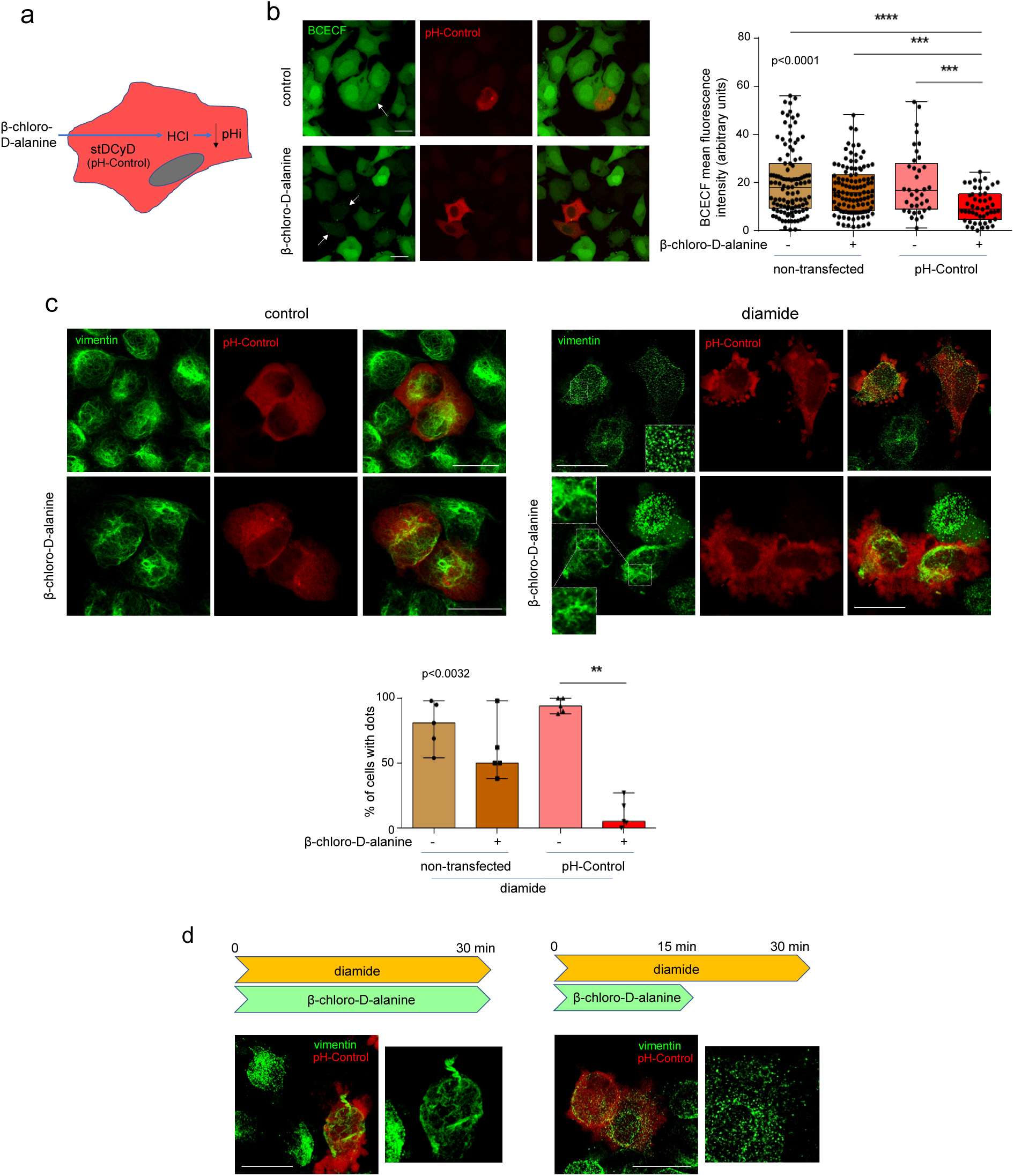
Intracellular acidification by a chemogenetic approach affords single cell and temporal modulation of diamide-elicited vimentin remodeling. (a) Scheme of the experimental approach based on transfection of cells with the pH-Control plasmid that encodes the bacterial enzyme StDCyD, which generates intracellular HCl, depending on the provision of the substrate, β-chloro-D-alanine. (b) For monitoring the effect of pH-Control on pHi, HeLa cells transfected with pH-Control (red), were loaded with the pH sensitive probe BCECF as detailed in Methods, and incubated in the absence or presence of 10 mM β-chloro-D-alanine. Image acquisition from live cells was started 2 min later. The mean fluorescence intensity from 4 experiments totaling at least 40 cells (between 40 and 114 cells) is shown in the box plot, where the central mark of the boxes is the median, the edges of the box are 25^th^ and 75^th^ percentiles, and whiskers extend to the most extreme data points. Statistical significance was assessed with the Kruskal-Wallis test (p value shown on the graph), and post hoc Dunńs multiple comparisons test, ***p<0.001, ****p<0.0001. (c) As indicated, HeLa cells transfected with pH-Control were incubated with 10 mM β-chloro-D-alanine for 2 min before the addition of 1 mM diamide for 15 min. At the end of the treatment cells were fixed and vimentin was detected by immunofluorescence. Insets display enlarged ROI. Images shown are representative of 5 independent experiments, from which the proportion of cells presenting vimentin in dots was calculated and is displayed in the box plot, where the top of the box is the median, and whiskers extend between maximum and minimum values. Statistical significance was assessed with the Kruskal-Wallis test (p value shown on the graph), and post hoc Dunńs multiple comparisons test, **p<0.01. The total number of cells monitored ranged from 93 to 697. (d) The reversibility of the protective effect of β-chloro-D-alanine was explored by following the incubation schemes outlined in the colored arrows, namely, keeping the coincubation of β-chloro-D-alanine and diamide for 30 min (left), or removing β-chloro-D-alanine for the last 15 min of the 30 min incubation with diamide (right). Cells were processed as above for detection of vimentin. Images at the right show enlarged views of the filamentous or dotted structures. Results are representative from 3 experiments. All images shown are overall projections. Scale bars, 20 µm.

Next, we explored the impact of genetic intracellular alkalinization. Overexpression of the Na+/H+ antiporter NHE1 has been reported to increase pHi in several cell types [56] (Fig. 7a). HeLa cells transfected with YFP-NHE1 displayed partial localization of the antiporter at the plasma membrane, together with an increase in pHi, indicated by the decrease in pHrodo fluorescence with respect to non-transfected cells (Fig. 7b). Remarkably, YFP-NHE1 expression also elicited a morphological change leading to an elongated fibroblast-like phenotype with cellular extensions (Fig. 7c, inset), illustrated by a significant decrease in the circularity of HNE1-expressing cells (Fig. 7c, graph). In non-transfected cells, treatment with diamide elicited a vimentin remodeling into dots, coexisting with some diffuse protein, filament remnants and condensations (Fig. 7d). In contrast, in numerous NHE1-transfected cells (>60%), diamide treatment led to the subcortical accumulation of vimentin positive structures (Fig. 7d, arrow).

**Figure 7.**
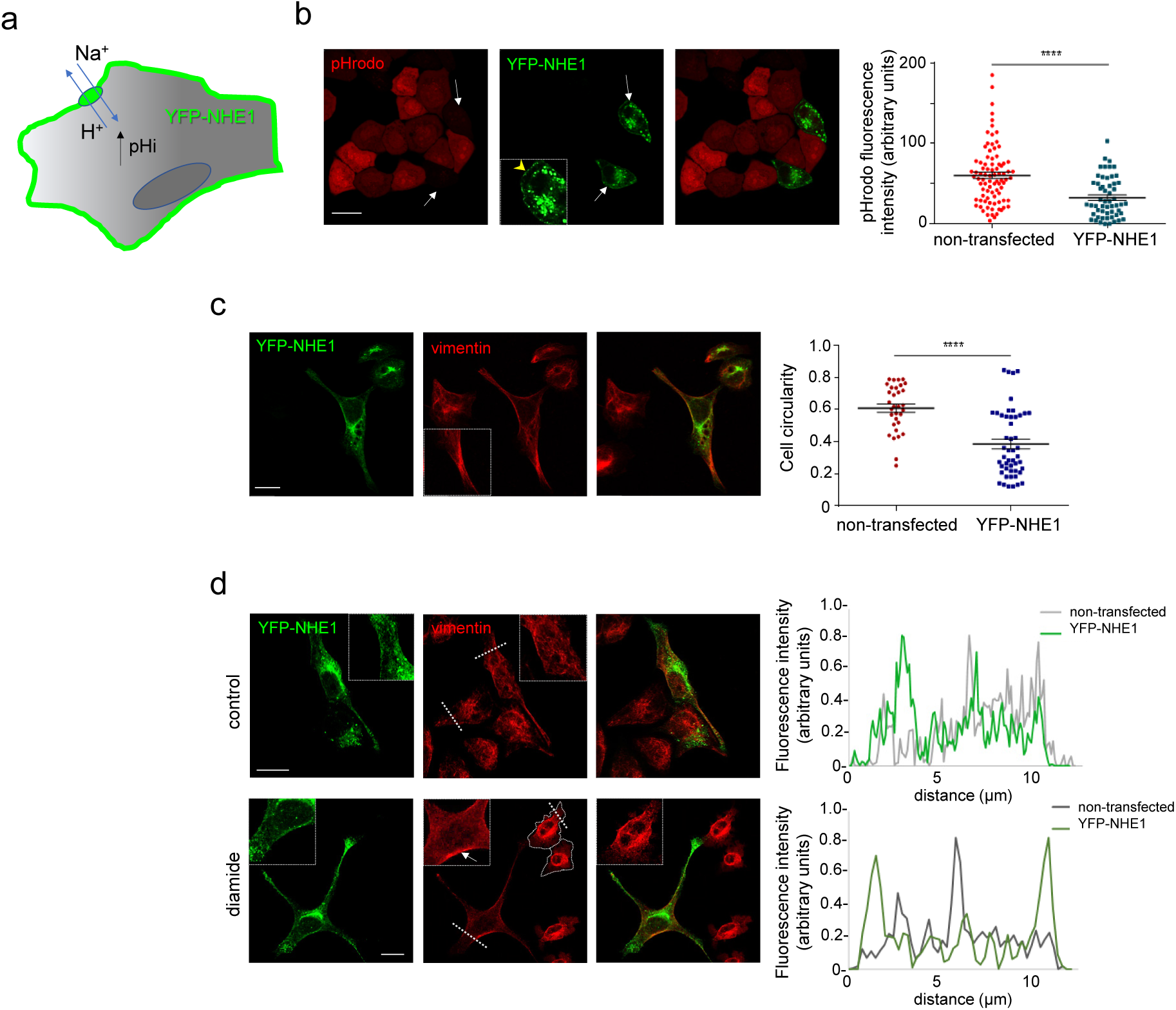
Intracellular alkalinization by transfection of NHE1 modulates diamide-elicited vimentin remodeling. (a) Schematic experimental approach. (b) The effect of YFP-NHE1 on pHi in HeLa cells was monitored with the pH sensitive probe pHrodo. Arrows point to YFP-NHE1 positive cells, with nearly undetectable pHrodo fluorescence. The arrowhead in the inset of the middle panel points to YFP-NHE1 lining the cell periphery. The fluorescence intensity of pHrodo from 5 independent experiments totaling 86 and 55 cells, respectively, is represented in the graph, which also displays average values ± SEM. Statistical significance was assessed with the Mann-Whitney test. ****p<0.0001. (c) HeLa cells were transfected with YFP-NHE1 and the circularity of non-transfected and transfected cells with detectable YFP-NHE1 at the cell membrane was measured. The graph shows determinations from at least 5 experiments totaling more than 27 cells, together with average values ± SEM. ****p<0.0001 by Student’s t-test. (d) HeLa cells transfected with YFP-HNE1 were treated with 1 mM diamide for 15 min, as indicated, and vimentin distribution was assessed by confocal microscopy. Insets show enlarged areas of interest. The peripheral accumulation of vimentin in diamide treated cells is indicated by the arrow in the inset, in which fluorescence intensity has been increased for better visualization. Fluorescence intensity profiles obtained along the dotted lines drawn on the images illustrate vimentin distribution in non-transfected (grey lines) and transfected (green lines) cells. The contour of several YFP-NHE1 negative cells has been highlighted. Note the presence of vimentin at the cell periphery after diamide treatment in YFP-NHE1 positive but not in negative cells. All images are overall projections. Scale bars, 20 µm in all panels.

This differential reorganization in transfected cells, illustrated in the fluorescence intensity profiles, (Fig. 7d, graphs), suggests a more intense vimentin remodeling by oxidative stress in cells expressing NHE1.

On the other hand, targeted induction of pHi alkalinization was achieved by transfection of the light gated outward proton pump ArchT (archaerhodopsin from Halorubrum strain TP009) [51]. As reported in other cell types [51, 52], and schematized in Fig. 8a, ArchT-BFP localized to the plasma membrane in HeLa cells (Fig. 8b). Photoactivation elicited a pHi increase, monitored with mCherry-SEpHluorin, significantly more intense in ArchT-BFP positive cells (Fig. 8b, graph). It is known that laser stimulation during confocal microscopy can elicit the generation of reactive oxygen species (ROS) [58]. Indeed, the photoactivation protocol increased the fluorescence of the ROS indicator DCF, more intensely in irradiated cells than in neighboring cells (Fig. 8c). Monitoring vimentin organization by cotransfection with a tracer amount of GFP-vimentin wt, which integrated into the endogenous network as previously reported [3, 12, 14] (Fig. 8d), revealed that photoactivation did not apparently perturb vimentin distribution in ArchT-BFP negative cells, and elicited relatively mild changes in their morphology consisting in appearance of blebs (Fig. 8d, upper images). In sharp contrast, photoactivation of cells expressing ArchT-BFP correlated with a marked retraction of vimentin filaments from the cell periphery, with condensation at the central region of the cell (Fig. 8d, lower images). In addition, the peripheral area devoid of filaments showed signs of vimentin disassembly, with appearance of diffuse protein, along with some fragments or dots (Fig. 8d, lower images, enlarged ROI). These alterations could result from increased pHi together with ROS generation. Strikingly, photoactivation elicited pronounced plasma membrane changes selectively in cells expressing ArchT-BFP, in which blebs coalesced giving rise to broad lamella (Fig. 8d, lower images). The proportion of cells undergoing membrane remodeling is shown in Fig. 8d, graph. A representative sequence of events is provided in Fig. 8e. These changes did not occur upon irradiation with light of a non-effective wavelength, namely, 488 nm (Supplementary Figure 7).

**Fig. 8.**
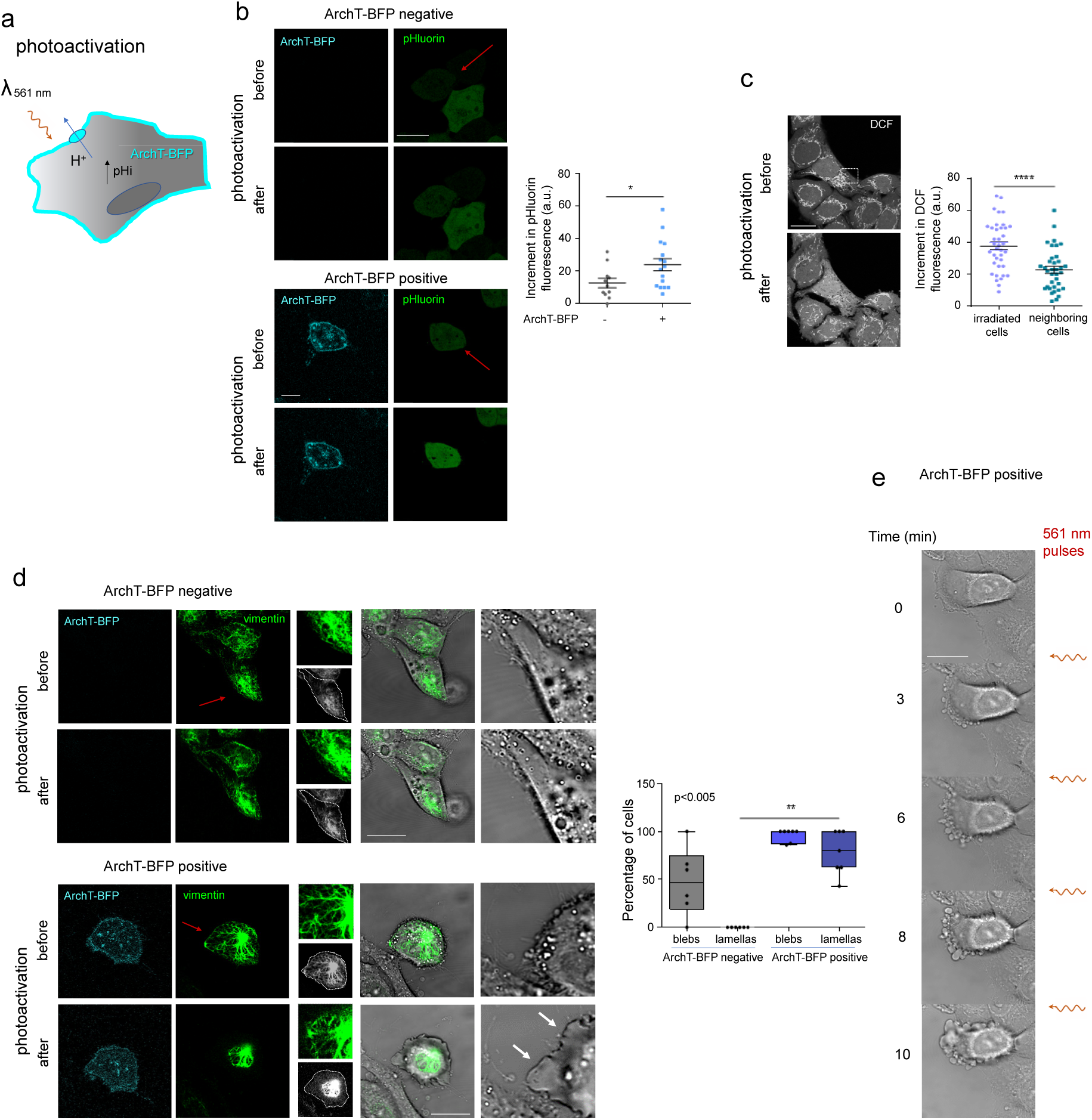
Intracellular alkalinization by an optogenetic approach induces single cell modulation of vimentin and membrane remodeling. (a) Scheme of the experimental approach based on cell transfection of ArchT-BFP, encoding a photoactivatable outward proton pump. (b) For monitoring the effect of ArchT-BFP photoactivation on pHi, HeLa cells were cotransfected with ArchT-BFP and mCherry-SEpHluorin, as indicated, and the cells marked by red arrows were subjected to the photoactivation protocol, as detailed in Methods. The graph depicts the increases in green fluorescence, taken as an index of the increase in pHi, measured in 3 independent experiments, totaling 16 ArchT-BFP positive cells and 11 negative cells (a.u., arbitrary units), and the average values ± SEM. *p<0.05 by Student’s t-test. (c) The generation of reactive oxygen species induced by the photoactivation protocol in the irradiated (dotted square) and the neighboring cells, was measured with the fluorescent probe DCF. The increases in DCF fluorescence from 4 experiments totaling 38 irradiated and 36 neighboring cells are shown in the graph, along the average values ± SEM. *p<0.0001 by Student’s t-test. (d) The remodeling of vimentin and the plasma membrane induced by the photoactivation protocol in ArchT-BFP negative (upper images) and positive (lower images) cells are shown. Photoactivated cells are marked by red arrows. Insets show details of vimentin filaments (upper image) and the position of the vimentin network with respect to the cell contour denoted by a dotted line (lower image, grayscale). Images on the right show the superimposition of vimentin and bright field images, and details of the membrane remodeling. Broad lamella are indicated by white arrows. The proportion of ArchT-BFP negative (non-transfected) or ArchT-BFP positive cells developing blebs or lamellas from at least six independent expriments is shown in the box plot at right. The top of each box is the median value, and the whiskers represent the difference between maximum and minimum values. Statistical analysis was assessed with the Kruskal-Wallis test (p value shown on the graph), followed by post hoc Dunńs multiple comparisons test,**p<0.01. (e) Representative sequence of events showing membrane remodeling of an ArchT-BFP positive cell at the beginning of the experiment and after each of 4 photoactivation steps, indicated by wavy arrows. Fluorescence images shown are single sections. All scale bars are 20 µm.

Taken together these results show that modulation of pHi, achieved by genetic, chemogenetic or optogenetic approaches, determines the response of vimentin to oxidative stress.

### EIPA attenuates vimentin disassembly at cell protrusions

We next explored whether vimentin disassembly could be modulated in association with physiological pH fluctuations. During cell migration, the leading edge undergoes a NHE1 mediated pH alkalinization that plays a key role in actin dynamics [39, 41] (scheme in Fig. 9a). Also, vimentin disassembles at the leading edge of migrating cells, and this is necessary for lamellipodia formation [59, 60]. Thus, we investigated a potential connection between leading edge alkalinization and vimentin disassembly by monitoring lamellipodia and vimentin in these structures in MEF wt in the absence or presence of EIPA to block NHE1. In control cells, vimentin filaments were absent from the leading edge or appeared disassembled into sparse squiggles or small particles, whereas EIPA treated cells showed more continuous vimentin structures close to the cell periphery (Fig. 9b). Hence, the distance between the migrating edge and the beginning of a robust continuous vimentin network was significantly shorter in EIPA treated cells (Fig. 9c). These differences were also evident when lamellipodia formation was stimulated by serum addition after starvation (Fig. 9b, c). Consistent with the restrictive role of vimentin filaments on lamellipodia formation reported previously [61], EIPA treated cells showed a blunted appearance with less prominent/smaller lamellipodia (quantitated in Fig. 9d). Similar observations were made under normal growth conditions and in serum stimulated cells (Fig. 9d). To assess a potential role of C328 in this process, we transfected MEF *Vim(-/-)* with vimentin wt or the C328H mutant, and stimulated lamellipodia formation by serum addition (Fig. 9e). We observed that, whereas vimentin wt filaments showed extensive disassembly into small particles at the cell periphery, intact vimentin C328H filaments frequently extended nearer the cell edge (quantitated in Fig. 9f).

**Fig. 9.**
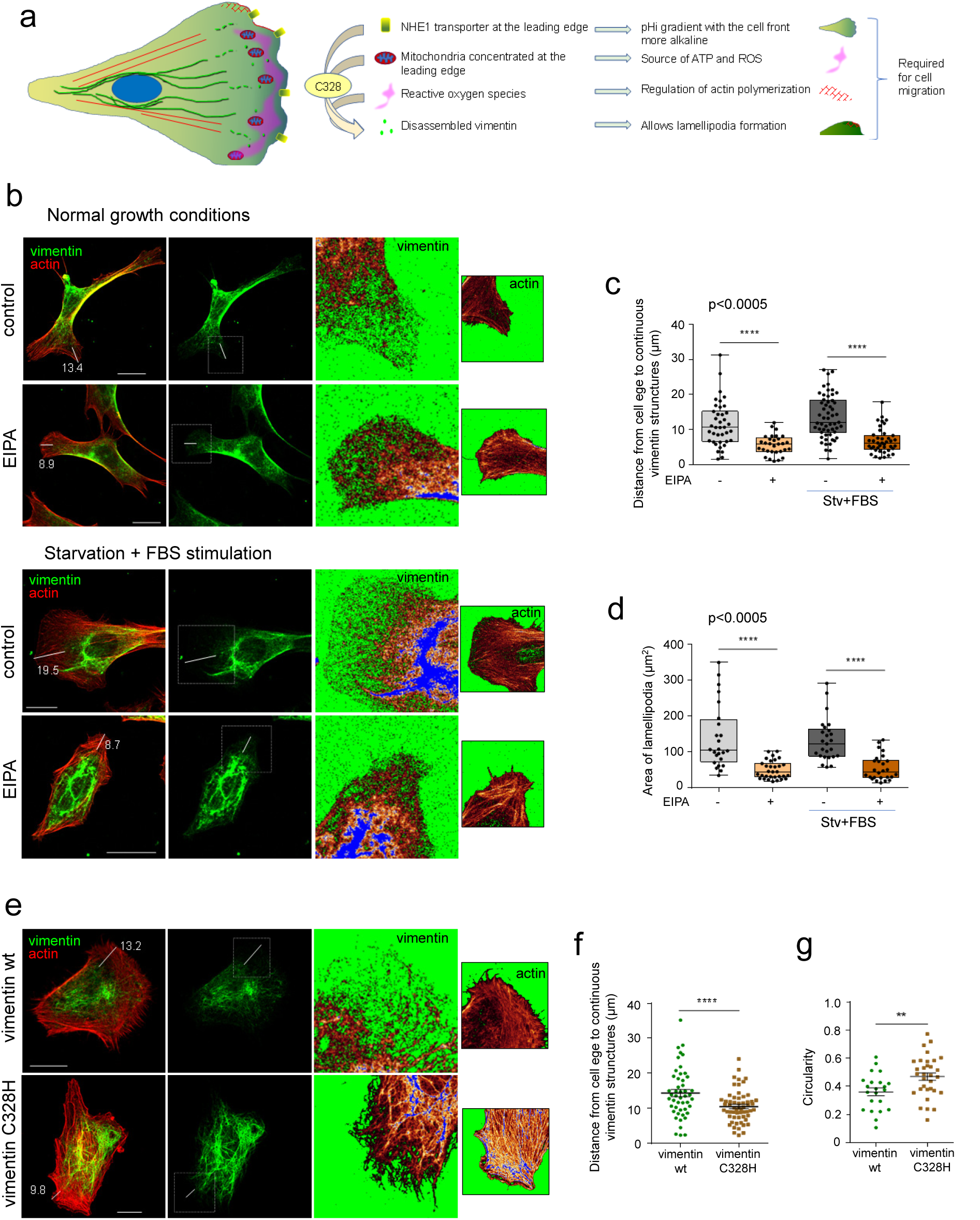
Importance of pHi in vimentin disassembly at cell protrusions. (a) Scheme of events reported to occur at the cell front during cell migration. These include generation of a NHE1-dependent pH gradient with alkalinization at the leading edge, production of ROS by mitochondria near the cell front, polymerization of actin in a reticular pattern, and disassembly of vimentin filaments, which is required for lamellipodia formation (see text for references). According to our hypothesis, blockade of NHE1 by EIPA would decrease vimentin susceptibility to ROS-elicited disassembly, and putatively, lamellipodia formation. In addition, mutation of C328, which represents the main pHi and redox responsive residue, would attenuate vimentin disassembly, negatively affecting lamellipodia. (b) MEF wt were monitored under normal culture conditions or after 20 h in 0.5% FBS (starvation, Stv) followed by 20 min serum re-addition to stimulate lamellipodia formation, as indicated. When specified, cells were incubated with 50 µM EIPA for the last 20 min of the incubation. Vimentin was detected by immunofluorescence and actin by Phalloidin-Alexa Fluor 488 or 568 staining. Distances between cell edges, delimited by actin staining, and the beginning of the denser vimentin filament network in single sections was measured in the segments displayed, the length of which is shown in µm. Images at the right show enlarged views of the corresponding ROI of the vimentin network (single sections) or actin (overall projections) displayed in the LUT mode for enhanced contrast of peripheral structures. Lamellipodia were identified by their typical morphology in the actin staining and their area was measured. Images shown are representative from 4 experiments. (c, d) Distances between the denser vimentin network and the cell edge (c) and area of lamellipodia (d) were monitored in 30-55 lamellipodia and 25-33 cells per experimental condition, respectively, and results are shown in the box plots, where the top of the boxes is the median, and the whiskers represent the difference between maximum and minimum values. Statistical significance was evaluated with the Kruskal-Wallis test (p value shown on the graphs), followed by post hoc Dunńs multiple comparisons test, ****p<0.0001. (e) MEF *Vim(-/-)* were transfected with vimentin wt or C328H, and lamellipodia formation was elicited by serum starvation followed by re-addition for 20 min. (f, g) Distances between the cell edge and the continuous vimentin network (f), and cell circularity (g), were monitored and calculated from three independent experiments, as described in Methods in at least 49 (f) and 22 (g) cells. Graphs depict individual and average values ± SEM. ****p<0.0001 by Student’s t-test. Scale bars, 20 µm.

Notably, cells expressing the C328H mutant often displayed a more compact morphology, with less cellular protrusions, which is reflected in their increased circularity (Fig. 9g, see Supplementary Figure 8 for additional examples), and the presence of lamellipodia was also less frequent (66±8% of cells expressing vimentin C328H showed lamellipodia, compared to 81.5% of vimentin wt expressing cells). In summary, these observations indicate that blocking NHE with EIPA decreases vimentin wt disassembly and lamellipodia formation. In addition, expression of a vimentin C328H mutant is associated with more intact filaments at the cell periphery and a lower proportion of cells with lamellipodia. Together, these results suggest that pHi and the presence of C328 influence vimentin disassembly at the leading edge, and that these factors may impact lamellipodia formation.

## Discussion

Vimentin is a key target of oxidative and electrophilic stress [18]. Its single cysteine residue, C328, can be modified by endogenous oxidants and electrophiles, natural compounds and drug metabolites [3, 12, 13, 17]. The relatively low pKa of C328 makes it susceptible to modification under pathophysiological conditions, as well as potentially sensitive to pH changes (schematized in Fig. 1a). Here we show that indeed, variations in pH modulate C328 modification. Acidic pH conditions, which would decrease the proportion of the C328 thiol in its reactive thiolate form, protect vimentin from modification and reorganization by oxidants, whereas alkaline pH conditions, which would favor the thiolate form, amplify their effects. This modulation can be achieved at the single cell level and in a spatiotemporal dependent manner. These observations unveil a new level of regulation of vimentin by redox alterations.

Cysteine residues are key elements in redox signaling [20]. Their high nucleophilicity enables the formation of covalent adducts with electrophiles and makes them preferred sites for a variety of PTMs. Cysteine reactivity towards specific agents is determined by their accessibility, involvement in hydrophobic interactions or formation of complexes, and pKa [62, 63]. In turn, pKa depends on their environment, particularly on the proximity of protonated carboxylic residues and/or hydrogen bonds donors [23, 64]. Cysteines with low pKa are typically present in redox regulated proteins, including thioredoxin, glutaredoxin, protein tyrosine phosphatases, etc. (reviewed in [23]). Examples of the modulation of cysteine susceptibility to oxidation or alkylation by pH variations in vitro, include the oxidation of protein tyrosine phosphatases [24], and the inactivation of glutaredoxin by iodoacetamide [25]. Our results show that vimentin C328 modification by oxidants and cysteine-reactive compounds is indeed modulated by pH changes from 7.0 to 8.0 in vitro, with alkaline pH favoring a more extensive modification and disruption of filamentous structures.

pHi can experiment ample changes depending on the subcellular compartment (from 4.7 in lysosomes to 7.8 in mitochondria), cell cycle, exposure to mitogens or growth factor deprivation (6.8 in certain forms of apoptosis) [38, 39, 41, 53, 65]. Although the reactivity of cysteine residues in cells is widely assumed to depend on pHi, examples of this modulation in the cellular context are scarce. Here we show that the effect of oxidants and electrophiles on cellular vimentin is modulated by pHi variations in the pathophysiological range. This modulation occurs in several cell types upon pHi variation by several strategies, including ionophores and pharmacological tackling of proton transporters. Thus, under acidic pHi conditions, vimentin filaments are protected from disruption by agents known to target C328 (diamide, H_2_O_2_ and HNE [3, 12, 13, 26]), whereas alkaline pHi potentiates their deleterious effect. Nevertheless, it should be noted that pHi fluctuations can exert cellular effects per se, including alteration of mitochondrial respiration, [34] or protein synthesis [65], modifications or interactions [66], for which the contribution of multiple, direct or indirect mechanisms to the modulation of vimentin disruption by pHi variations cannot be excluded. In spite of this, the fact that vimentin mutants lacking C328 are protected from remodeling by diamide, even at high pH, supports the importance of this residue in the sensitizing effect of alkaline pH.

Interestingly, strategies reflecting the free thiol content of vimentin after the various treatments indicate that diamide, alone or in combination with monensin, decreases free thiol accessibility, whereas when diamide treatment is performed at acidic pHi, i.e., in presence of EIPA, C328 thiol availability tends to be more similar to control conditions. Nevertheless, it should be noted that cysteine modifications can be structurally very diverse and, in some cases, progressive or interchangeable. Moreover, a certain degree vimentin C328 occupancy can occur under control conditions [13, 67], and shuttling of the modifications or conformational changes with exposure of unmodified C328 can also occur [67]. In turn, diamide can promote both reversible and irreversible cysteine modifications, including different degrees of oxidation, with potentially different functional consequences. Additionally, in the context of the filament, modifications could display cooperativity or competition due to conformational alterations [68, 69]. Therefore, the possibilities for modulation at this level are multiple and highly complex.

Most oxidants, including diamide and H_2_O_2_, also alter actin structures and microtubules [3, 70]. Interestingly, the protective effect of acidic pHi herein observed appears selective for vimentin since neither actin nor tubulin networks were protected from diamide-elicited disruption by lowering pHi. This postulates vimentin filaments, and likely other type III intermediate filaments, namely GFAP, as cytoskeletal proteins especially susceptible to the modulation of the modification of their single cysteine residue by pHi changes. Nevertheless, the actin cytoskeleton is considered one of the main pH-sensitive systems in cells [66]. Several actin regulatory proteins are regulated by pHi. For instance, cofilin, which elicits actin depolymerization, is more active at alkaline pH [71], whereas talin binding to f-actin decreases at pH above 7.2, allowing fast focal adhesion turnover and cell migration (reviewed in [66]). In these cases, the protonation state of key histidine residues is critical [30, 72]. However, to the best of our knowledge, there is no information on the pH regulation of actin dynamics involving the susceptibility of cysteine residues to modification.

Nonetheless, pHi changes could affect the propensity to oxidation of other cellular proteins. For instance, kinases including PKA, PKG and Aurora [73, 74], and phosphatases, such as PP2A, PP2B and various PTP are regulated by redox mechanisms (reviewed in [75]). Therefore, remodeling of the vimentin network by certain oxidants could involve the combined action of oxidative modifications and changes in phosphorylation. Indeed, we previously observed that treatment of cells with calyculin A, an inhibitor of protein phosphatases 1 and 2A, increases vimentin phosphorylation at S72 and the proportion of soluble vimentin, and potentiates diamide elicited disruption [12], whereas elimination of 11 phosphorylation sites in the vimentin N-terminal domain confers partial protection [12]. In addition, other proteins, including chaperones and cytolinkers may be pH sensitive and contribute to the effects observed. The observation that vimentin C328H and C328A mutants are not sensitive to diamide-elicited remodeling, even at alkaline pHi, highlights the importance of a thiol group in this position for the response of the protein. However, the influence of pHi changes on other parameters, such as vimentin charge and stability [76], mechanics [77], or electrostatic interactions between subunits within filaments, and/or with cations (e.g. Zn^2+^, Mg^2+^) [26], cannot be discarded.

Importantly, we have demonstrated that modulation of pHi by chemogenetic or optogenetic strategies affords spatiotemporal regulation of vimentin remodeling by oxidants. Indeed, selective intracellular acidification with pH-Control elicits a reversible protection of vimentin against disruption by diamide, depending on the presence of the substrate β-Cl-D-Ala. Strategies eliciting intracellular alkalinization exert the opposite effect, potentiating vimentin disruption and/or altering its pattern. Thus, transfection of NHE1 alkalinizes cells and impacts vimentin redistribution upon diamide treatment. Moreover, intracellular alkalinization achieved by photoactivation of the proton pump ArchT, elicits vimentin disassembly and/or retraction from the cell periphery selectively in ArchT-expressing photoactivated cells. This associates with a remarkable membrane remodeling with blebbing and formation of lamellipodia-like cell protrusions. Interestingly, membrane blebbing has been reported to occur preferentially at cell edges lacking abundant vimentin filaments [78]. Moreover, vimentin depletion has been shown to increase blebbing, and specifically elicit the formation of leader blebs, involved in ameboid cell migration [79]. Generation of ROS, occurring upon laser stimulation (Fig. 8c), could probably contribute to these effects.

Indeed, cell migration is a physiological process dependent on the cooperation of pHi and ROS gradients. The relatively alkaline pH of the cell front during migration [41], together with the local generation of ROS [80], could favor the disassembly of vimentin that occurs at the leading edge, which is required for lamellipodia formation [60]. Here we have observed that inhibition of intracellular alkalinization by EIPA attenuates vimentin disassembly at the cell periphery in MEF expressing vimentin wt and blunts cell protrusions. Moreover, filaments of vimentin C328H, which is resistant to disruption under all pHi conditions, extend closer to the cell edges than those of vimentin wt, in association with less abundant lamellipodia in MEF *Vim(-/-)* transfected with this construct. Cytoplasmic alkalinization at the leading edge also plays a key role in actomyosin contractility driving cell migration through the pH dependent modulation of actin regulatory proteins [41]. We and others have previously reported a negative effect of vimentin filaments on actomyosin contractility that is relieved by vimentin disassembly [13, 81]. Therefore, it could be hypothesized that pH and C328 dependent vimentin disassembly may favor actin driven cell protrusions.

The regulation of vimentin disassembly by the combined action of oxidants and pHi variations could have important implications in health and disease. For instance, the higher pHi of tumoral cells [28] could favor higher vimentin dynamics in response to redox changes, or sensitize the vimentin cytoskeleton to pharmacological disruption. In a general cellular context, vimentin could provide a second level of response to oxidative or electrophilic stress. In contrast to proteins bearing cysteine residues with a very low pKa, which are expected to sense reactive species in a wide pH range, a large proportion of vimentin would be well suited to mediate a response in the physiological and moderately alkaline pHi ranges, hypothetically related to general or local rises in pHi, such as those occurring during cell division or migration.

In summary, our results reinforce the role of the single, conserved cysteine residue of vimentin in filament and network organization, and propose that it may concomitantly respond to redox status and pHi variations in a spatiotemporal manner, thus contributing to the complex mechanisms of vimentin regulation in cells. Moreover, our results provide a relevant demonstration of the modulation of cysteine-mediated redox responses by pHi variations.

## Data availability statement

All data generated in this study are provided in the Supplementary Information or available from the authors.

## Funding

This work was supported by Grants RTI2018-097624-B-I00 and PID2021-126827OB-I00, funded by MCIN /AEI/10.13039/501100011033 and ERDF, A way of making Europe; “Astromad”, ref. LCF/PR/HR21/52410002, from Fundación “la Caixa”; AEM is the recipient of a Juan de la Cierva postdoctoral contract FJC2021-047028-I funded by MCIN/AEI/ 10.13039/501100011033 and European Union NextGenerationEU/PRTR; PGJ is the recipient of a predoctoral contract PRE2019-088194, from MCIN/AEI /10.13039/501100011033 and ESF, Investing in your future”, CVV has been the recipient of a fellowship JAEIntroICU-2021-LifeHub-03, from the network PIE-202120E047-Conexiones-Life (CSIC), Spain.

## Abbreviations

DCF, 2′,7′-dichlorofluorescein diacetate; EIPA, 5-(N-ethyl-N-isopropyl)amiloride; GFAP, glial fibrillary acidic protein; HNE, 4-hydroxynonenal; Mal-B, Maleimide-PEG_2_-Biotin; pHi, intracellular pH; PTM, posttranslational modification(s); ROI, region of interest; TRITC, tetramethylrhodamine B isothiocyanate.

## Acknowledgements

Feedback from EpiLipidNet (CA19105) is gratefully acknowledged. We thank the personnel of the Optical Microscopy facility at Centro de Investigaciones Biológicas Margarita Salas for their advice. We are indebted to Dr. Asal Ghaffari Zaki and Prof. Emrah Eroglu (Istanbul Medipol University, Turkiye) for sharing the pH-Control plasmid.

## Author contributions

A.E.M., C.V.V. and D.P.S. performed the research and analyzed the data. P.M.C. and P.G.J. performed research. A.E.M., M.A.P. and D.P.S. wrote the paper. D.P.S. designed and supervised the study, obtained and managed grants. All authors reviewed and edited the paper.

## Notes

### Competing Interest Statement

The authors have declared no competing interest.

### Summary of Updates

V2 contains new results and analyses

## References

[1] A.E. Patteson, A. Vahabikashi, R.D. Goldman, P.A. Janmey, Mechanical and Non-Mechanical Functions of Filamentous and Non-Filamentous Vimentin, Bioessays 42(11) (2020) e2000078.

[2] A.B. Ndiaye, G.H. Koenderink, M. Shemesh, Intermediate Filaments in Cellular Mechanoresponsiveness: Mediating Cytoskeletal Crosstalk From Membrane to Nucleus and Back, Frontiers in cell and developmental biology 10 (2022) 882037.

[3] D. Pérez-Sala, C.L. Oeste, A.E. Martínez, B. Garzón, M.J. Carrasco, F.J. Cañada, Vimentin filament organization and stress sensing depend on its single cysteine residue and zinc binding, Nature communications 6 (2015) 7287.

[4] P. Mohanasundaram, L.S. Coelho-Rato, M.K. Modi, M. Urbanska, F. Lautenschlager, F. Cheng, J.E. Eriksson, Cytoskeletal vimentin regulates cell size and autophagy through mTORC1 signaling, PLoS Biol 20(9) (2022) e3001737.

[5] K.M. Ridge, J.E. Eriksson, M. Pekny, R.D. Goldman, Roles of vimentin in health and disease, Genes Dev 36(7-8) (2022) 391–407.

[6] N. Mucke, T. Wedig, A. Burer, L.N. Marekov, P.M. Steinert, J. Langowski, U. Aebi, H. Herrmann, Molecular and biophysical characterization of assembly-starter units of human vimentin, J Mol Biol 340(1) (2004) 97–114.

[7] H. Herrmann, U. Aebi, Intermediate Filaments: Structure and Assembly, Cold Spring Harbor perspectives in biology 8(11) (2016) a018242.

[8] S.A. Eldirany, I.B. Lomakin, M. Ho, C.G. Bunick, Recent insight into intermediate filament structure, Curr Opin Cell Biol 68 (2021) 132–143.

[9] M. Eibauer, M.S. Weber, R. Kronenberg-Tenga, C.T. Beales, R. Boujemaa-Paterski, Y. Turgay, S. Sivagurunathan, J. Kraxner, S. Köster, R.D. Goldman, O. Medalia, Vimentin filaments integrate low complexity domains in a highly complex helical structure, Nat Struct Mol Biol (2024) 31:939–949.

[10] G. Colakoglu, A. Brown, Intermediate filaments exchange subunits along their length and elongate by end-to-end annealing, J Cell Biol 185(5) (2009) 769–77.

[11] F. Nunes Vicente, M. Lelek, J.Y. Tinevez, Q.D. Tran, G. Pehau-Arnaudet, C. Zimmer, S. Etienne-Manneville, G. Giannone, C. Leduc, Molecular organization and mechanics of single vimentin filaments revealed by super-resolution imaging, Sci Adv 8(8) (2022) eabm2696.

[12] A. Mónico, S. Duarte, M.A. Pajares, D. Pérez-Sala, Vimentin disruption by lipoxidation and electrophiles: role of the cysteine residue and filament dynamics, Redox biology 23 (2019) 101098.

[13] P. González-Jiménez, S. Duarte, A. Martínez-Fernández, E. Navarro-Carrasco, V. Lalioti, M.A. Pajares, D. Pérez-Sala, Vimentin single cysteine residue acts as a tunable sensor for network organization and as a key for actin remodeling in response to oxidants and electrophiles, Redox biology 64 (2023) 102756.

[14] P. Martínez-Cenalmor, A.E. Martínez, D. Moneo-Corcuera, P. González-Jiménez, D. Pérez-Sala, Oxidative stress elicits the remodeling of vimentin filaments into biomolecular condensates, Redox biology 75 (2024) 103282.

[15] K. Stamatakis, F.J. Sánchez-Gómez, D. Pérez-Sala, Identification of novel protein targets for modification by 15-deoxy-D12,14-prostaglandin J2 in mesangial cells reveals multiple interactions with the cytoskeleton., J Am Soc Nephrol 17 (2006) 89–98.

[16] D. Frescas, C.M. Roux, S. Aygun-Sunar, A.S. Gleiberman, P. Krasnov, O.V. Kurnasov, E. Strom, L.P. Virtuoso, M. Wrobel, A.L. Osterman, M.P. Antoch, V. Mett, O.B. Chernova, A.V. Gudkov, Senescent cells expose and secrete an oxidized form of membrane-bound vimentin as revealed by a natural polyreactive antibody, Proc Natl Acad Sci U S A 114(9) (2017) E1668–E1677.

[17] M. Kaus-Drobek, N. Mucke, R.H. Szczepanowski, T. Wedig, M. Czarnocki-Cieciura, M. Polakowska, H. Herrmann, A. Wyslouch-Cieszynska, M. Dadlez, Vimentin S-glutathionylation at Cys328 inhibits filament elongation and induces severing of mature filaments in vitro, FEBS J 287 (2020) 5304–5322.

[18] A. Viedma-Poyatos, M.A. Pajares, D. Pérez-Sala, Type III intermediate filaments as targets and effectors of electrophiles and oxidants, Redox biology 36 (2020) 101582.

[19] D. Pérez-Sala, R. Quinlan, The redox-responsive roles of intermediate filaments in cellular stress detection, integration and mitigation, Curr Opin Cell Biol 86 (2024) 102283.

[20] J.M. Held, Redox Systems Biology: Harnessing the Sentinels of the Cysteine Redoxome, Antioxid Redox Signal 32(10) (2020) 659–676.

[21] H. Sies, D.P. Jones, Reactive oxygen species (ROS) as pleiotropic physiological signalling agents, Nat Rev Mol Cell Biol 21(7) (2020) 363–383.

[22] M.J. Davies, Protein oxidation and peroxidation, Biochem J 473(7) (2016) 805–25.

[23] G. Roos, N. Foloppe, J. Messens, Understanding the pK(a) of redox cysteines: the key role of hydrogen bonding, Antioxid Redox Signal 18(1) (2013) 94–127.

[24] A. Groen, S. Lemeer, T. van der Wijk, J. Overvoorde, A.J. Heck, A. Ostman, D. Barford, M. Slijper, J. den Hertog, Differential oxidation of protein-tyrosine phosphatases, J Biol Chem 280 (2005) 10298–10304.

[25] S.C. Jao, S.M. English Ospina, A.J. Berdis, D.W. Starke, C.B. Post, J.J. Mieyal, Computational and mutational analysis of human glutaredoxin (thioltransferase): probing the molecular basis of the low pKa of cysteine 22 and its role in catalysis, Biochemistry 45(15) (2006) 4785–96.

[26] A. Mónico, J. Guzman-Caldentey, M.A. Pajares, S. Martin-Santamaria, D. Pérez-Sala, Molecular Insight into the Regulation of Vimentin by Cysteine Modifications and Zinc Binding, Antioxidants (Basel) 10(7) (2021) 1039.

[27] S.B. Wall, J.Y. Oh, A.R. Diers, A. Landar, Oxidative modification of proteins: an emerging mechanism of cell signaling, Frontiers in physiology 3 (2012) 369.

[28] K.A. White, B.K. Grillo-Hill, D.L. Barber, Cancer cell behaviors mediated by dysregulated pH dynamics at a glance, J Cell Sci 130(4) (2017) 663–669.

[29] E. Persi, M. Duran-Frigola, M. Damaghi, W.R. Roush, P. Aloy, J.L. Cleveland, R.J. Gillies, E. Ruppin, Systems analysis of intracellular pH vulnerabilities for cancer therapy, Nature communications 9(1) (2018) 2997.

[30] Y. Liu, K.A. White, D.L. Barber, Intracellular pH Regulates Cancer and Stem Cell Behaviors: A Protein Dynamics Perspective, Front Oncol 10 (2020) 1401.

[31] W.H. Moolenaar, Effects of growth factors on intracellular pH regulation, Annu Rev Physiol 48 (1986) 363–76.

[32] S.J. Reshkin, A. Bellizzi, S. Caldeira, V. Albarani, I. Malanchi, M. Poignee, M. Alunni-Fabbroni, V. Casavola, M. Tommasino, Na+/H+ exchanger-dependent intracellular alkalinization is an early event in malignant transformation and plays an essential role in the development of subsequent transformation-associated phenotypes, FASEB J 14(14) (2000) 2185–97.

[33] A. Rebollo, J. Gómez, A.M.d. Aragón, P. Lastres, A. Silva, D. Pérez-Sala, Apoptosis Induced by IL-2 Withdrawal is Associated with an Intracellular Acidification, Experimental Cell Research 218 (1995) 581–585.

[34] A.J. Genders, S.D. Martin, S.L. McGee, D.J. Bishop, A physiological drop in pH decreases mitochondrial respiration, and HDAC and Akt signaling, in L6 myocytes, Am J Physiol Cell Physiol 316(3) (2019) C404–C414.

[35] G.J. Gores, A.L. Nieminen, B.E. Wray, B. Herman, J.J. Lemasters, Intracellular pH during “chemical hypoxia” in cultured rat hepatocytes. Protection by intracellular acidosis against the onset of cell death, J Clin Invest 83(2) (1989) 386–96.

[36] H. Hagberg, Intracellular pH during ischemia in skeletal muscle: relationship to membrane potential, extracellular pH, tissue lactic acid and ATP, Pflugers Arch 404(4) (1985) 342–7.

[37] M.J. Yong, B. Kang, U. Yang, S.S. Oh, J.H. Je, Live Streaming of a Single Cell’s Life over a Local pH-Monitoring Nanowire Waveguide, Nano Lett 22(15) (2022) 6375–6382.

[38] J.S. Spear, K.A. White, Single-cell intracellular pH dynamics regulate the cell cycle by timing the G1 exit and G2 transition, J Cell Sci 136(10) (2023) jcs260458.

[39] J.R. Casey, S. Grinstein, J. Orlowski, Sensors and regulators of intracellular pH, Nat Rev Mol Cell Biol 11(1) (2010) 50–61.

[40] J.J. Rennick, C.J. Nowell, C.W. Pouton, A.P.R. Johnston, Resolving subcellular pH with a quantitative fluorescent lifetime biosensor, Nature communications 13(1) (2022) 6023.

[41] C. Martin, S.F. Pedersen, A. Schwab, C. Stock, Intracellular pH gradients in migrating cells, Am J Physiol Cell Physiol 300(3) (2011) C490–5.

[42] H. Herrmann, I. Hofmann, W.W. Franke, Identification of a nonapeptide motif in the vimentin head domain involved in intermediate filament assembly, J Mol Biol 223(3) (1992) 637–50.

[43] A. Mónico, E. Rodríguez-Senra, F.J. Cañada, S. Zorrilla, D. Pérez-Sala, Drawbacks of dialysis procedures for removal of EDTA, PLoS ONE 12(1) (2017) e0169843.

[44] A. Krezel, W. Maret, The biological inorganic chemistry of zinc ions, Arch Biochem Biophys 611 (2016) 3–19.

[45] S. Kirchhof, A. Strasser, H.J. Wittmann, V. Messmann, N. Hammer, A.M. Goepferich, F.P. Brandl, New insights into the cross-linking and degradation mechanism of Diels-Alder hydrogels, J Mater Chem B 3(3) (2015) 449–457.

[46] E.R. Hondorp, R.G. Matthews, Oxidation of cysteine 645 of cobalamin-independent methionine synthase causes a methionine limitation in Escherichia coli, J Bacteriol 191(10) (2009) 3407–10.

[47] A.J. Sarria, J.G. Lieber, S.K. Nordeen, R.M. Evans, The presence or absence of a vimentin-type intermediate filament network affects the shape of the nucleus in human SW-13 cells, J Cell Sci 107 (Pt 6) (1994) 1593–607.

[48] D. Moneo-Corcuera, A. Viedma-Poyatos, K. Stamatakis, D. Pérez-Sala, Desmin Reorganization by Stimuli Inducing Oxidative Stress and Electrophiles: Role of Its Single Cysteine Residue, Antioxidants (Basel) 12(9) (2023) 1703.

[49] M. Koivusalo, C. Welch, H. Hayashi, C.C. Scott, M. Kim, T. Alexander, N. Touret, K.M. Hahn, S. Grinstein, Amiloride inhibits macropinocytosis by lowering submembranous pH and preventing Rac1 and Cdc42 signaling, J Cell Biol 188(4) (2010) 547–63.

[50] A. Ghaffari Zaki, S.M. Miri, S. Cimen, T. Akgul Caglar, E.N. Yigit, M.S. Aydin, G. Ozturk, E. Eroglu, Development of a Chemogenetic Approach to Manipulate Intracellular pH, J Am Chem Soc 145(22) (2023) 11899–11902.

[51] L. Wu, A. Dong, L. Dong, S.Q. Wang, Y. Li, PARIS, an optogenetic method for functionally mapping gap junctions, eLife (2019) 14:8:e43366.

[52] C.E.T. Donahue, M.D. Siroky, K.A. White, An Optogenetic Tool to Raise Intracellular pH in Single Cells and Drive Localized Membrane Dynamics, J Am Chem Soc 143(45) (2021) 18877–18887.

[53] D. Pérez-Sala, D. Collado-Escobar, F. Mollinedo, Intracellular Alkalinization Suppresses Lovastatin-induced Apoptosis in HL-60 Cells through the Inactivation of a pH-dependent Endonuclease, J. Biol. Chem. 270(11) (1995) 6235–6242.

[54] C.Y. Yang, P.W. Chang, W.H. Hsu, H.C. Chang, C.L. Chen, C.C. Lai, W.T. Chiu, H.C. Chen, Src and SHP2 coordinately regulate the dynamics and organization of vimentin filaments during cell migration, Oncogene 38(21) (2019) 4075–4094.

[55] P. Vigne, C. Frelin, E.J. Cragoe, Jr., M. Lazdunski, Ethylisopropyl-amiloride: a new and highly potent derivative of amiloride for the inhibition of the Na+/H+ exchange system in various cell types, Biochem Biophys Res Commun 116(1) (1983) 86–90.

[56] C.H. Man, F.E. Mercier, N. Liu, W. Dong, G. Stephanopoulos, L. Jiang, Y. Jung, C.P. Lin, A.Y.H. Leung, D.T. Scadden, Proton export alkalinizes intracellular pH and reprograms carbon metabolism to drive normal and malignant cell growth, Blood 139(4) (2022) 502–522.

[57] A.V. Berezhnov, M.P. Soutar, E.I. Fedotova, M.S. Frolova, H. Plun-Favreau, V.P. Zinchenko, A.Y. Abramov, Intracellular pH Modulates Autophagy and Mitophagy, J Biol Chem 291(16) (2016) 8701–8.

[58] E. Alexandratou, D. Yova, P. Handris, D. Kletsas, S. Loukas, Human fibroblast alterations induced by low power laser irradiation at the single cell level using confocal microscopy, Photochem Photobiol Sci 1(8) (2002) 547–52.

[59] B.T. Helfand, M.G. Mendez, S.N. Murthy, D.K. Shumaker, B. Grin, S. Mahammad, U. Aebi, T. Wedig, Y.I. Wu, K.M. Hahn, M. Inagaki, H. Herrmann, R.D. Goldman, Vimentin organization modulates the formation of lamellipodia, Mol Biol Cell 22(8) (2011) 1274–89.

[60] M.E. Kidd, D.K. Shumaker, K.M. Ridge, The role of vimentin intermediate filaments in the progression of lung cancer, Am J Respir Cell Mol Biol 50(1) (2014) 1–6.

[61] R.A. Battaglia, S. Delic, H. Herrmann, N.T. Snider, Vimentin on the move: new developments in cell migration, F1000Research 7 (2018) 1796.

[62] G. Ferrer-Sueta, B. Manta, H. Botti, R. Radi, M. Trujillo, A. Denicola, Factors affecting protein thiol reactivity and specificity in peroxide reduction, Chem Res Toxicol 24(4) (2011) 434–50.

[63] G. Gambardella, G. Cattani, A. Bocedi, G. Ricci, New Factors Enhancing the Reactivity of Cysteines in Molten Globule-Like Structures, International journal of molecular sciences 21(18) (2020) 6949.

[64] L.B. Poole, The basics of thiols and cysteines in redox biology and chemistry, Free Radic Biol Med 80 (2015) 148–57.

[65] M. Flinck, S.H. Kramer, S.F. Pedersen, Roles of pH in control of cell proliferation, Acta Physiol (Oxf) 223(3) (2018) e13068.

[66] M. Damaghi, J.W. Wojtkowiak, R.J. Gillies, pH sensing and regulation in cancer, Frontiers in physiology 4 (2013) 370.

[67] E. Griesser, V. Vemula, A. Mónico, D. Pérez-Sala, M. Fedorova, Dynamic posttranslational modifications of cytoskeletal proteins unveil hot spots under nitroxidative stress, Redox biology 44 (2021) 102014.

[68] L.M. Koharudin, H. Liu, R. Di Maio, R.B. Kodali, S.H. Graham, A.M. Gronenborn, Cyclopentenone prostaglandin-induced unfolding and aggregation of the Parkinson disease-associated UCH-L1, Proc Natl Acad Sci U S A 107(15) (2010) 6835–40.

[69] C.A. Lemmon, T. Ohashi, H.P. Erickson, Probing the folded state of fibronectin type III domains in stretched fibrils by measuring buried cysteine accessibility, J Biol Chem 286(30) (2011) 26375–82.

[70] J. Huot, F. Houle, F. Marceau, J. Landry, Oxidative stress-induced actin reorganization mediated by the p38 mitogen-activated protein kinase/heat shock protein 27 pathway in vascular endothelial cells, Circ Res 80(3) (1997) 383–92.

[71] N. Yonezawa, E. Nishida, H. Sakai, pH control of actin polymerization by cofilin, J Biol Chem 260(27) (1985) 14410–2.

[72] J. Srivastava, G. Barreiro, S. Groscurth, A.R. Gingras, B.T. Goult, D.R. Critchley, M.J. Kelly, M.P. Jacobson, D.L. Barber, Structural model and functional significance of pH-dependent talin-actin binding for focal adhesion remodeling, Proc Natl Acad Sci U S A 105(38) (2008) 14436–41.

[73] F. Cuello, P. Eaton, Cysteine-Based Redox Sensing and Its Role in Signaling by Cyclic Nucleotide-Dependent Kinases in the Cardiovascular System, Annu Rev Physiol 81 (2019) 63–87.

[74] D.P. Byrne, S. Shrestha, M. Galler, M. Cao, L.A. Daly, A.E. Campbell, C.E. Eyers, E.A. Veal, N. Kannan, P.A. Eyers, Aurora A regulation by reversible cysteine oxidation reveals evolutionarily conserved redox control of Ser/Thr protein kinase activity, Science signaling 13(639) (2020) eaax2713.

[75] J. den Hertog, A. Groen, T. van der Wijk, Redox regulation of protein-tyrosine phosphatases, Arch Biochem Biophys 434 (2005) 11–15.

[76] B.A. Unger, C.Y. Wu, A.A. Choi, C. He, K. Xu, Hypersensitivity of the vimentin cytoskeleton to net-charge states and Coulomb repulsion, bioRxiv (2024) doi: 10.1101/2024.07.08.602555.

[77] A.V. Schepers, C. Lorenz, S. Koster, Tuning intermediate filament mechanics by variation of pH and ion charges, Nanoscale 12(28) (2020) 15236–15245.

[78] A.S. Chikina, A.O. Zholudeva, M.E. Lomakina, Kireev, II, A.A. Dayal, A.A. Minin, M. Maurin, T.M. Svitkina, A.Y. Alexandrova, Plasma Membrane Blebbing Is Controlled by Subcellular Distribution of Vimentin Intermediate Filaments, Cells 13(1) (2024) 105.

[79] S.B. Lavenus, S.M. Tudor, M.F. Ullo, K.W. Vosatka, J.S. Logue, A flexible network of vimentin intermediate filaments promotes migration of amoeboid cancer cells through confined environments, J Biol Chem 295(19) (2020) 6700–6709.

[80] V.V. Pak, D. Ezerina, O.G. Lyublinskaya, B. Pedre, P.A. Tyurin-Kuzmin, N.M. Mishina, M. Thauvin, D. Young, K. Wahni, S.A. Martinez Gache, A.D. Demidovich, Y.G. Ermakova, Y.D. Maslova, A.G. Shokhina, E. Eroglu, D.S. Bilan, I. Bogeski, T. Michel, S. Vriz, J. Messens, V.V. Belousov, Ultrasensitive Genetically Encoded Indicator for Hydrogen Peroxide Identifies Roles for the Oxidant in Cell Migration and Mitochondrial Function, Cell Metab 31(3) (2020) 642–653 e6.

[81] Y. Jiu, J. Peranen, N. Schaible, F. Cheng, J.E. Eriksson, R. Krishnan, P. Lappalainen, Vimentin intermediate filaments control actin stress fiber assembly through GEF-H1 and RhoA, J Cell Sci 130(5) (2017) 892–902.

