## Supplementary material for "Intracellular pH modulates vimentin remodeling in response to oxidants"

Running title: Vimentin as a paradigm of the pH dependence of redox signaling

**Supplementary Figures**

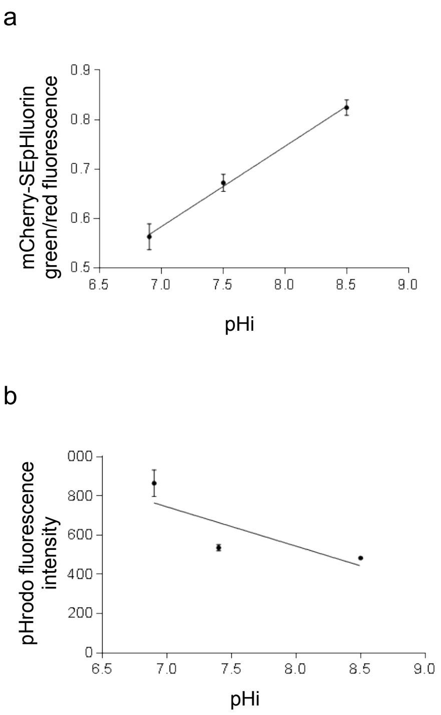


**Supplementary Figure 1. Examples of calibration assays showing the pH dependence of the fluorescence of mCherry-SEpHluorin and pHrodo.** (a) SW13/Cl.2 cells expressing mCherry-SEpHluorin were cultured in a $K^+$-rich buffer at the indicated extracellular pH in the presence of the ionophore nigericin, as detailed in Methods. The ratio of green to red fluorescence was calculated from eight fields from two experiments and is represented as average values ± SEM per every pH condition. (b) HeLa cells were loaded with pHrodo, as detailed in Methods, and cultured in LCIS medium at different pH, as above. The intensity of the red fluorescence for every indicated pH was quantitated from eight fields from two experiments and is shown as average values ± SEM.

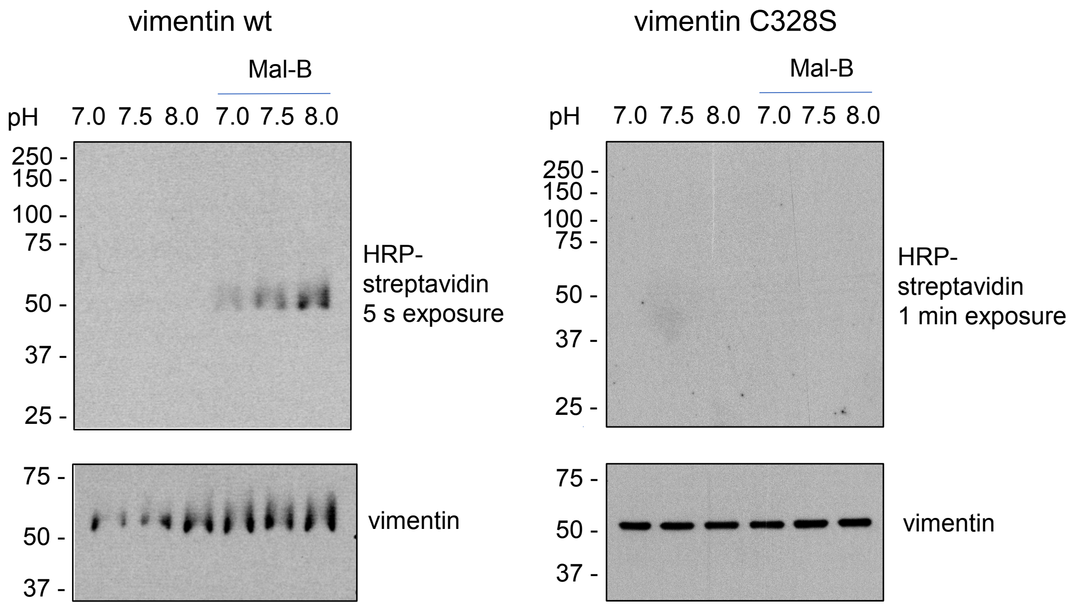


**Supplementary Figure 2: Lack of incorporation of Mal-B in a vimentin C328S mutant.** Purified recombinant vimentin wt or C328S at 4 µM was incubated with Mal-B for 15 min at different pH, and analyzed by reducing SDS-PAGE, as described in Fig. 1. The position of molecular weight markers, in kDa, is shown at the left. Incorporation of biotin into the purified protein was assessed by blot with HRP-streptavidin and ECL detection. Images show exposures of 5 s for vimentin wt and 1 min for vimentin C328S, under the same conditions than vimentin wt. Total vimentin in membranes was assessed by western blot.

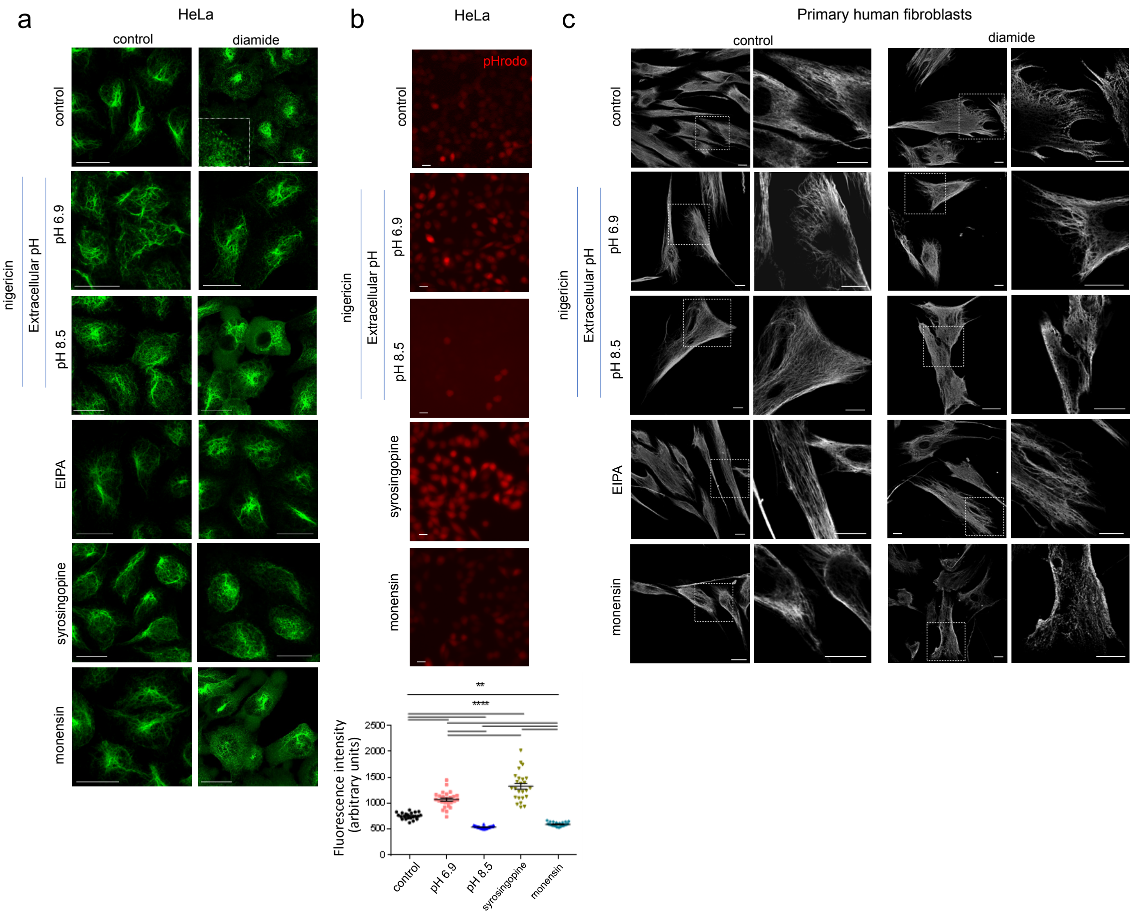


**Supplementary Figure 3. The disruption of the vimentin network elicited by diamide is modulated by pH in several cell types.** HeLa cells (a) or human primary fibroblasts (c) were incubated in culture media at various pH supplemented with nigericin as indicated, or pretreated with 25 µM EIPA for 15 min, 10 µM syrosingopine for 1 h or 10 µM monensin for 5 min, before treatment in the absence or presence of 1 mM diamide for 15 min. The distribution of vimentin was assessed by immunofluorescence (scale bars, 20 µm). (b) The pHi of HeLa cells achieved by the various strategies was monitored by incubation with pHrodo as detailed in the experimental section, before treatment in culture media at various pH supplemented with nigericin for 15 min as indicated, treatment with 10 µM syrosingopine for 75 min, or with 10 µM monensin for 20 min. Afterwards, live cells were imaged. Two independent experiments were performed and at least 24 cells were evaluated. Data shown are mean values ± SEM. For comparisons of multiple data sets the one-way analysis of variance (ANOVA) followed by Tukey's multiple comparison test was used. Statistically significant differences are indicated on the graph, as follows: **p<0.01; ****p<0.0001. In (c), images on the left for each condition are overall projections. Enlarged views of areas of interest delimited by dotted squares are depicted at the right of each image, as single sections. Scale bars, 20 µm.

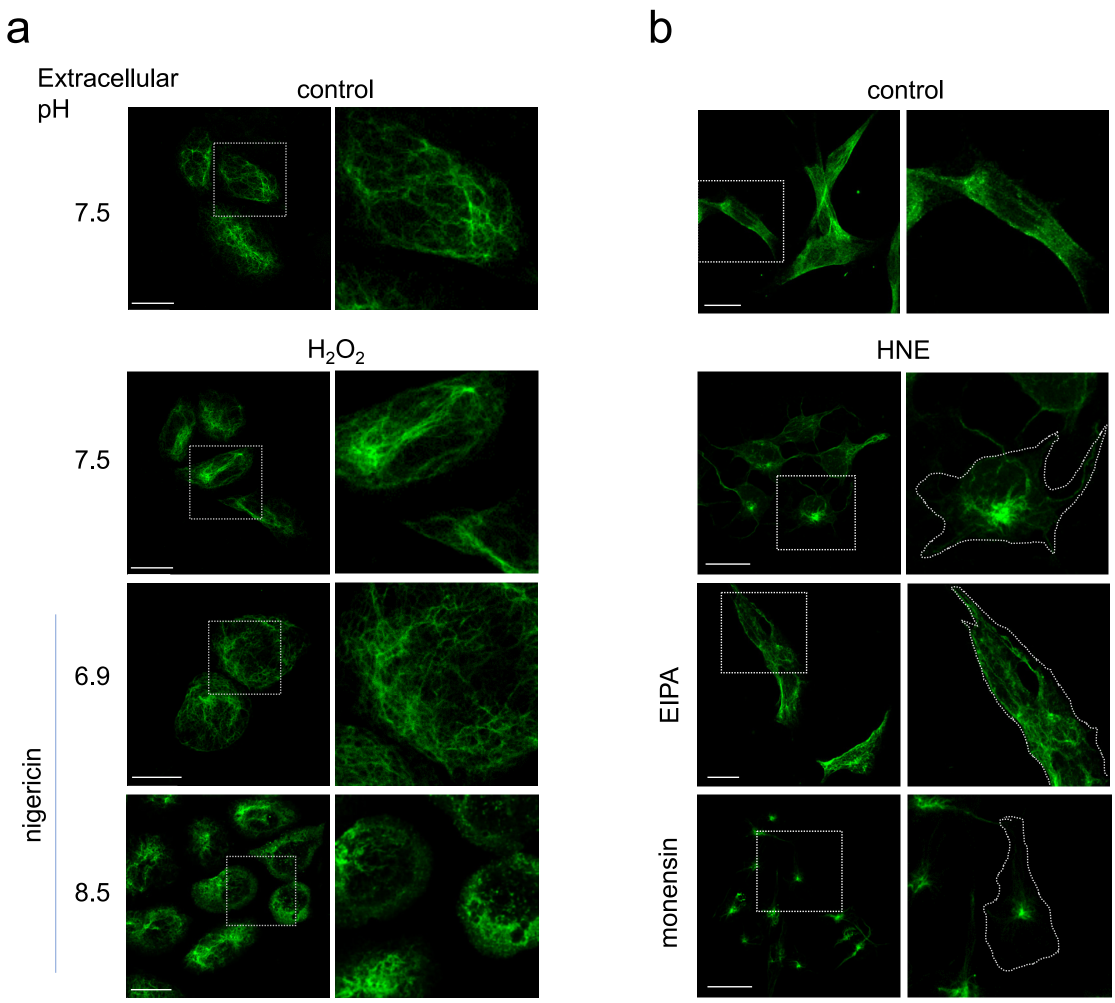


**Supplementary Figure 4. Modulation of the effects of $H_2O_2$ and HNE on the vimentin network by variations in pHi.** (a) SW13/cl.2 cells expressing vimentin wt were incubated in culture media at various pH supplemented with nigericin as indicated, and treated in the absence or presence or 1 mM $H_2O_2$ for 25 min. (b) MEF wt were treated with 10 µM HNE for 2 h alone or after preincubation with 25 µM EIPA for 15 min or 10 µM monensin for 5 min. The reorganization of vimentin was assessed by immunofluorescence. Enlarged views of areas of interest delimited by dotted squares are depicted at the right. Results are representative from three experiments. Scale bars, 20 µm.

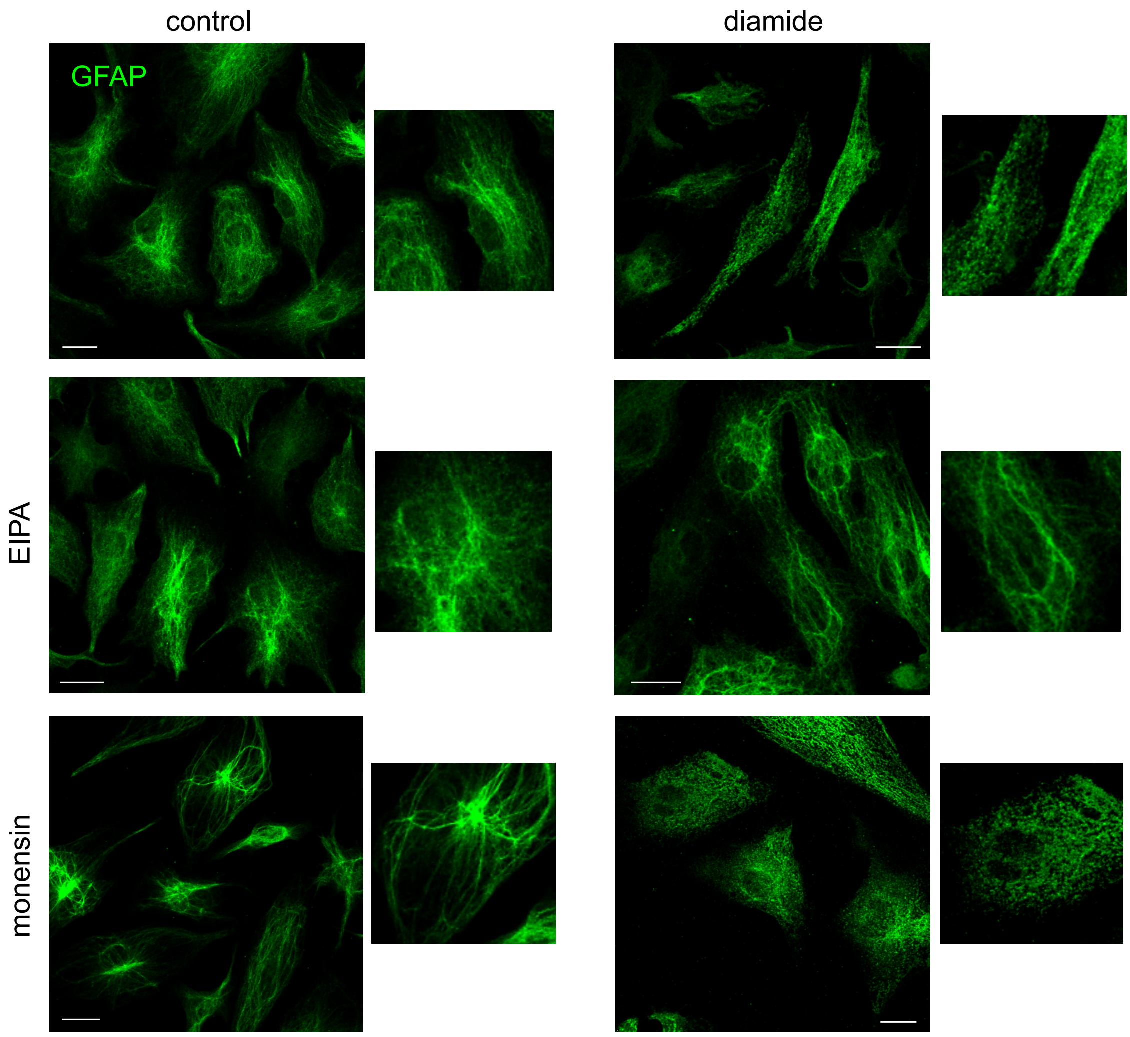


**Supplementary Figure 5. Modulation of diamide-elicited GFAP remodeling by pHi.** U-251 MG cells, which express endogenous GFAP were treated with 1 mM diamide for 15 min. As indicated, cells were pretreated with 25 µM EIPA for 15 min or 10 µM monensin for 5 min. After treatment, cells were fixed and processed for GFAP detection by immunofluorescence. Scale bars, 20 µm.

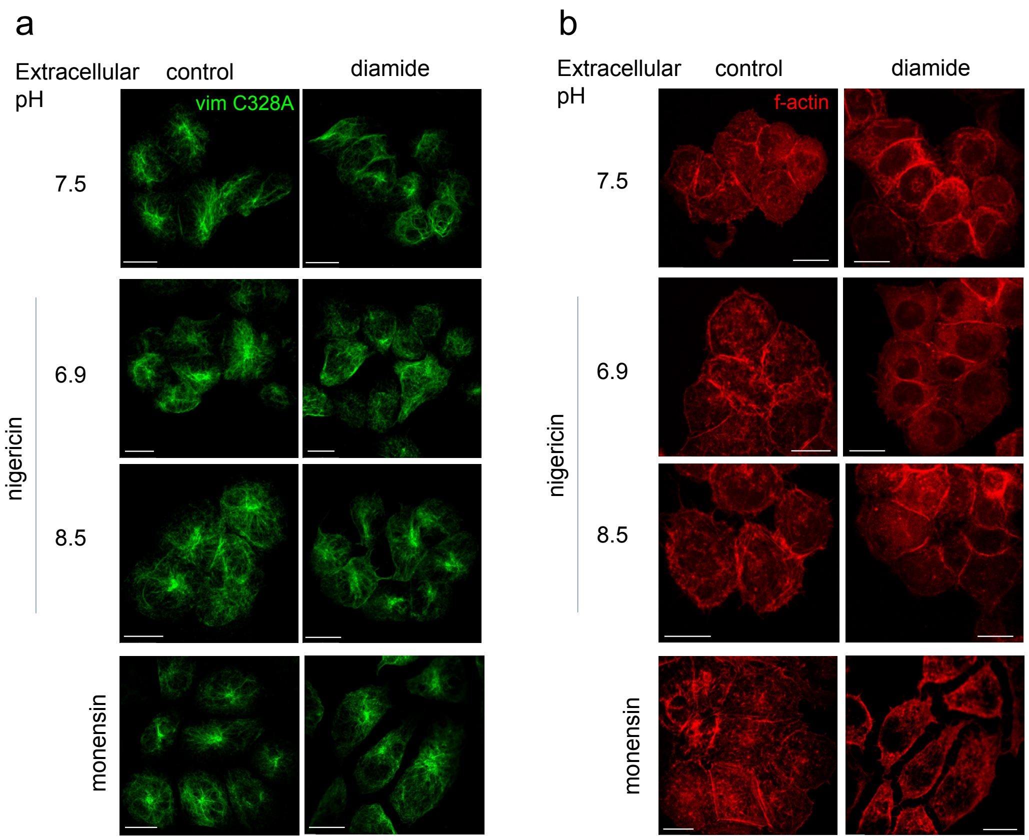


**Supplementary Figure 6. Importance of the vimentin single cysteine residue, C328, in the disruption by diamide at different pH.** SW13/cl.2 cells expressing the vimentin C328A mutant were incubated in culture media at different pH supplemented with nigericin as indicated, or pretreated with 10 µM monensin for 5 min, before treatment in the absence or presence of 1 mM diamide for 15 min. The distribution of vimentin filaments was assessed by immunofluorescence (a) and that of f-actin by staining with TRITC-phalloidin (b). Scale bars, 20 µm.

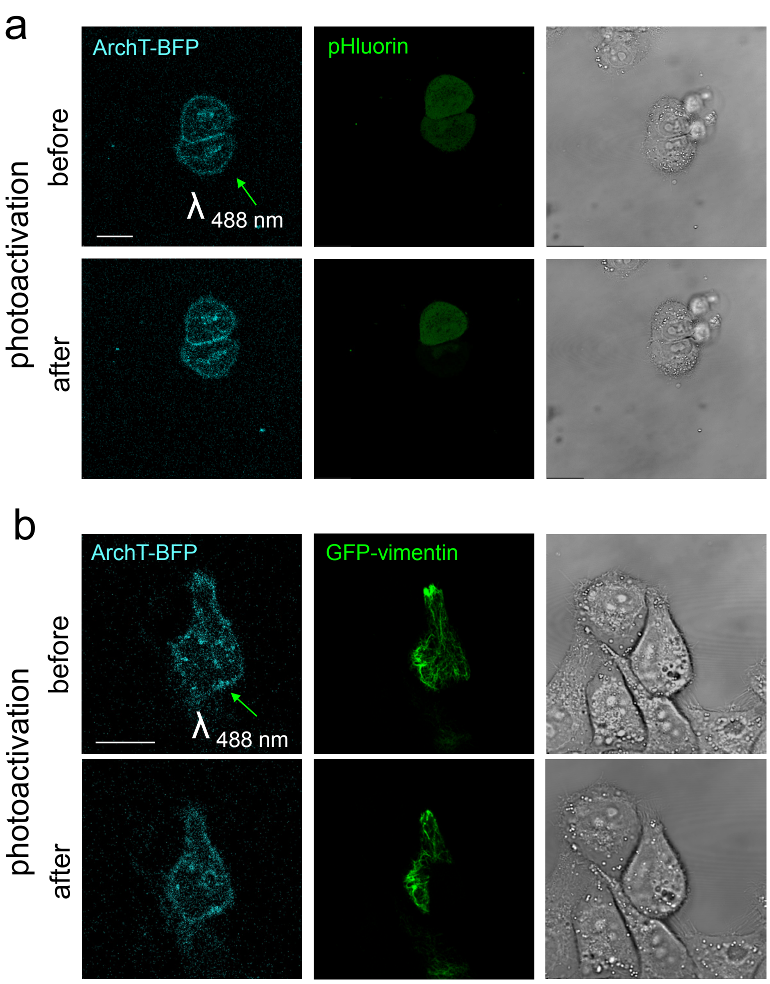


**Supplementary Figure 7. Photoactivation with 488 nm light does not induce plasma membrane or vimentin remodeling in cells expressing ArchT-BFP.** (a) HeLa cells were cotransfected with ArchT-BFP and mCherry-SEpHluorin, and one of the cells (marked by an arrow) was subjected to the photoactivation protocol, as detailed in Methods, using the 488 nm laser line. Note that this protocol completely bleaches SEpHluorin fluorescence in the photoactivated cell (center) and does not induce significant alterations of the plasma membrane, as illustrated in the bright field images (right). (b) Cells were cotransfected with Arch-T-BFP and GFP-vimentin, and the cell marked by the arrow was photoactivated as above. This protocol elicited bleaching of the GFP-vimentin signal in the irradiated region (center), without significantly perturbing the vimentin network or the plasma membrane (right). Scale bars, 20 µm.

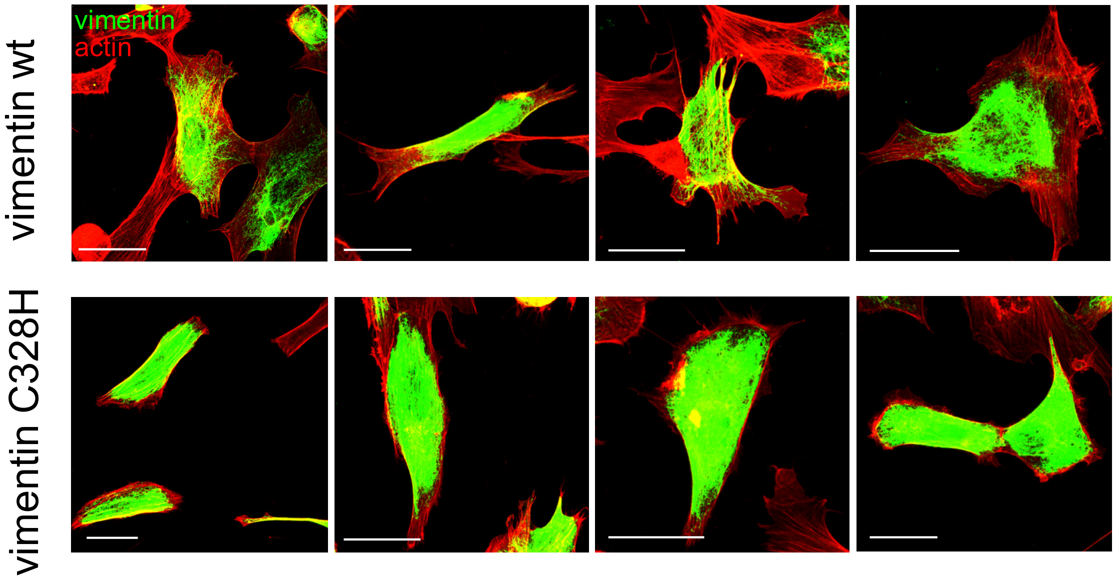


**Supplementary Figure 8. Morphology of MEF *Vim(-/-)* transfected with vimentin wt or C328H.** MEF *Vim(-/-)* were transfected with vimentin wt or C328H, as indicated, and the distribution of vimentin and a ctin was observed by confocal microscopy as detailed in Methods. Images shown provide several examples of the morphological features of MEF after 20 h in 0.5% (v/v) FBS (starvation) followed by 20 min serum re-addition to stimulate lamellipodia formation, as in Fig. 9e. All images were acquired under the same conditions. Please note that the signal of vimentin is deliberately overexposed in order to illustrate the disassembled structures present mainly at the cell periphery in MEF *Vim(-/-)* transfected with vimentin wt. Scale bars, 20 µm.
